# Synthetic Condensed-Phase Signaling Expands Kinase Specificity and Responds to Macromolecular Crowding

**DOI:** 10.1101/2021.12.10.472132

**Authors:** Dajun Sang, Tong Shu, Liam J. Holt

## Abstract

Liquid–liquid phase separation (LLPS) can concentrate biomolecules and accelerate reactions within membraneless organelles. For example, the nucleolus and PML-nuclear bodies are thought to create network hubs by bringing signaling molecules such as kinases and substrates together. However, the mechanisms and principles connecting mesoscale organization to signaling dynamics are difficult to dissect due to the pleiotropic effects associated with disrupting endogenous condensates. Here, we recruited multiple distinct kinases and substrates into synthetic LLPS systems to create new phosphorylation reactions within condensates, and generally found increased activity and broadened specificity. Dynamic phosphorylation within condensates could drive cell-cycle-dependent localization changes. Detailed comparison of phosphorylation of clients with varying recruitment valency and affinity into condensates comprised of either flexible or rigid scaffolds revealed unexpected principles. First, high client concentration within condensates is important, but is not the main factor for efficient multi-site phosphorylation. Rather, the availability of a large number of excess client binding sites, together with a flexible scaffold is crucial. Finally, phosphorylation within a suboptimal, flexible condensate was modulated by changes in macromolecular crowding. Thus, condensates readily generate new signaling connections and can create sensors that respond to perturbations to the biophysical properties of the cytoplasm.

## Introduction

Eukaryotic cells use functionally distinct compartments for spatiotemporal control of complex biochemical reactions. Organelles such as the endoplasmic reticulum or Golgi apparatus are bound by membranes, enabling the segregation of molecules using specific transport systems. On the other hand, there is increasing evidence for the importance of membraneless subcellular compartments, such as nucleoli, stress granules, germ (P) granules, Cajal bodies (Banani et al., 2017). These biomolecular condensates can be highly dynamic and sometimes form through liquid–liquid phase separation (LLPS) (Shin and Brangwynne, 2017). Recently, intensive research has revealed mechanisms of condensate formation, but it has been more difficult to determine the importance of LLPS for the modulation of biochemical reactions.

Recently, a number of studies have characterized biochemical reactions within condensates. For example, actin assembly is accelerated in the nephrin–NCK–N-WASP system (Li et al., 2012); SOS activation is increased in LAT-Grb2-SOS condensates (Huang et al., 2019); ribozyme catalysis and RNA polymerization is enhanced by coacervation (Poudyal et al., 2019). However, it has been difficult to determine the precise mechanisms underlying these increased reaction rates. The simplest hypothesis is that increased local concentrations accelerate reactions. However, additional mechanisms have been proposed. For example, LAT-Grb2-SOS condensates increased the membrane dwell time of key components, thus increasing SOS activation (Huang et al., 2019). In another study on SUMOylation, a decrease in apparent K_M_ was induced by scaffolding within condensates (Peeples and Rosen, 2021). However, it is difficult to understand the impact of condensation on biological regulation because mutations that disrupt LLPS or that perturb recruitment of clients to condensates can have pleiotropic effects. For example, while it is possible to mutate a protein sequence such that it fails to undergo LLPS, any associated loss of activity could be due to this loss of condensation or equally could be due to an unrelated loss of intrinsic protein function. A handful of excellent studies have surmounted this problem through orthogonal reconstitution of the condensation behavior. For example, Petry and colleagues demonstrated the importance of phase-separation *per se* in microtubule nucleation by reconstituting the activity and phase separation of microtubule nucleation factor TPX2 with an orthogonal disordered domain (King and Petry, 2020). However, there are very few examples, and it can be difficult to rescue endogenous biological functions in this way.

Synthetic biology is a powerful approach to investigate general principles while avoiding pleiotropic effects and unknown, evolved cellular regulation that can confound interpretation. Pioneering work has attempted to develop new functional systems within synthetic condensates that are orthogonal to the biology of the host cell. For example, Lemke and colleagues built orthogonal synthetic condensates to compartmentalize translation, allowing incorporation of distinct noncanonical amino acids specific target proteins (Reinkemeier et al., 2019; Reinkemeier and Lemke, 2021). Additionally, synthetic condensates have been used to study the biophysics of phase separation in living cells (Nakamura et al., 2019). For example, synthetic optogenetic systems, including ‘OptoDroplets’ and ‘Corelets’, were developed to exert spatiotemporal control of phase transitions in cells, permitting quantitative mapping of intracellular phase diagrams (Bracha et al., 2019; Shin et al., 2017). These systems can be broadly investigated without the constraints of endogenous biology and have been highly effective in revealing general principles of phase separation in cells.

The complex cellular environment can strongly affect LLPS. The cell interior is highly crowded, both in the cytosol and nucleus (Ellis and Minton, 2003b; Luby-Phelps, 1999; Luby-Phelps, 2013; van den Berg et al., 2017). There are multiple sources for intracellular crowding. Intracellular structures such as dynamic cytoskeletal networks in the cytosol (Trepat et al., 2008) and chromatin networks in the nucleus create confinement (Stephens et al., 2019). In addition, up to 40% of the cell volume is excluded by macromolecules that do not form inter-connected networks, including both proteins, ribosomes (Delarue et al., 2018), and nucleic acids (Ellis and Minton, 2003b; Luby-Phelps, 1999). The crowded cellular environment is thought to be close to a glass transition; indeed, sudden decreases in motion are observed at the mesoscale (tens of nanometers) when ATP is depleted from the cell (Parry et al., 2014; Zhou et al., 2009), suggesting that non-equilibrium active processes are required to fluidize the cell. Recent studies have focused on the impact of crowded molecular networks on LLPS. In the cytoplasm, mTORC1 modulates crowding by tuning ribosome concentration, and this has strong effects on phase separation (Delarue et al., 2018). In the nucleus, the chromatin network can mechanically suppress the coalescence and ripening of OptoDroplets, thus impacting condensate number, size and placement (Lee et al., 2021; Zhang et al., 2021b).

Similar phenomena have also been observed in engineered LLPS systems using synthetic polymer networks (Rosowski et al., 2020a; Rosowski et al., 2020b; Vidal-Henriquez and Zwicker, 2021). Therefore, it is clear that the crowded, active cell interior modulates phase separation. It follows, therefore, that condensates should be capable of sensing the cellular environment and transducing this information to chemical signals such as protein phosphorylation. However, there is no direct evidence of this phenomenon.

Protein kinases regulate most aspects of cellular activity, including cell growth, differentiation, proliferation, and apoptosis. Kinase activity is strictly regulated, and kinases must discriminate between numerous proteins to find appropriate substrates. Defects in kinase activity regulation or loss of specificity can lead to human diseases such as cancer (Cicenas et al., 2018). Kinases exploit multiple mechanisms to achieve specificity, including subcellular localization, docking motif interactions, and recognition of consensus phosphorylation sites (Alexander et al., 2011; Howard et al., 2014; Mok et al., 2010; Remenyi et al., 2006). For example, Mitogen Associated Protein Kinases (MAP kinases) and Cyclin Depedent Kinase (CDK) family kinases have the highest affinity for consensus phosphorylation motifs with a proline residue C-terminal to the phosphoacceptor site (Howard et al., 2014; Mok et al., 2010). These two kinase families also use docking motifs to selectively bind to and phosphorylate substrates (Faustova et al., 2021; Howard et al., 2014; Miller and Turk, 2018; Ord and Loog, 2019; Ord et al., 2019). Several recent studies showed that a large number of kinases reside in condensates (Wippich et al., 2013; Zhang et al., 2021a), but it remains poorly understood how kinase activity is modified by recruitment to these structures. Therefore, it is of great interest to understand how kinase reaction specificity and activity can be affected within condensates. Here we use synthetic biology approaches to begin to uncover principles of condensed-phase signaling. We recruited several kinases and substrates as clients into multiple types of synthetic condensates. These kinases included the MAP kinases, ERK1 and Fus3, and the Cyclin Dependent Kinase 1, Cdk1. We found that phosphorylation was increased by client recruitment into condensates both *in vitro* and *in vivo*. Kinase specificity was expanded in condensates indicating that condensation can facilitate the creation of new links in phosphoregulatory networks. Substrates were phosphorylated in the absence of a docking motif, and on non-consensus phosphoacceptor sequences. Phosphorylation within condensates could respond to dynamic changes in kinase activity within minutes. Systematic variation of scaffold and client properties revealed that, beyond increasing client concentration (mass action), the availability of excess client binding sites within the condensates and the flexibility of scaffolds strongly impacted reaction acceleration in condensates. Finally, we found that phosphorylation within condensates can respond to molecular crowding, thus creating a biophysical sensor.

## Results

### Recruitment to synthetic condensates increases phosphorylation rates *in vitro*

Initially, we engineered a system to investigate the effect of recruitment of a kinase and substrate to a synthetic condensate *in vitro*. Our synthetic phase separated condensates were based on multivalent interactions between a tandem repeat of ten Small Ubiquitin-like Modifier (SUMO) domains (SUMO_10_) and a second polypeptide with six repeats of a SUMO interacting motif (SIM_6_) (Figure 1A). SUMO_10_ and SIM_6_ readily undergo liquid-liquid phase-separation (LLPS) when mixed (Banani et al., 2016). Throughout this paper, we refer to proteins that form condensates as “scaffolds”, and call proteins that are recruited to synthetic condensates “clients”.

**Figure 1:**
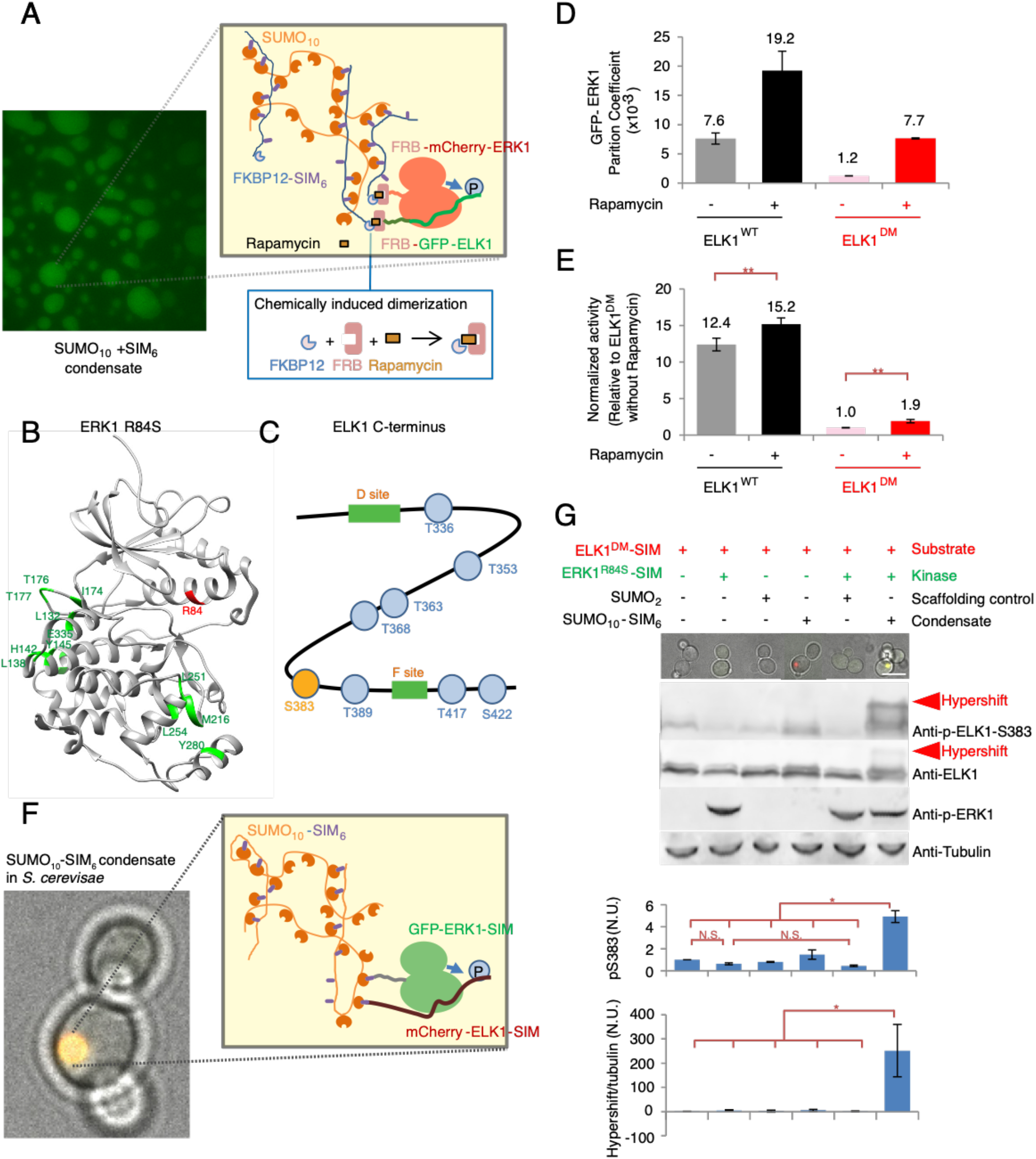
Recruitment into synthetic condensates increases phosphorylation rates. (A) Schematic showing the chemical dimerization approach used to rapidly recruit clients into SUMO_10_ + SIM_6_ condensates *in vitro*. Mixture of purified recombinant protein SUMO_10_ and SIM_6_ leads to condensate formation. The rapamycin-induced dimerization system was used to recruit the kinase ERK1 and substrate ELK1 clients into condensates. SIM_6_ was fused with FKBP12, and clients were fused with FRB. Rapamycin induces dimerization of FKBP12 with FRB. (B) Structure of constitutively active ERK1^R84S^ (PDB ERK1): R84 shown in red; surfaces that interact with ELK1 docking-motifs are shown in green. (C) Structure of the ELK1 C-terminus substrate. Known ERK1/2 phosphorylation sites are shown as blue circles, the phosphoepitope at S383 is highlighted orange, and the docking motifs colored in green. (D) ELK1 can be chemically recruited to condensates. The partition coefficient of GFP-ELK1 in SUMO_10_ + SIM_6_ condensates was quantified with and without addition of rapamycin. Error bars indicate ± SD, n = 2. (E) Recruitment to condensates increases phosphorylation rates *in vitro*. Quantification of kinase assays measuring ELK1 phosphorylation rates with and without rapamycin treatment. The C-termini of ELK1 (ELK1^WT^) or an ELK1 docking motif mutant (ELK1^DM^) were used as substrates. Error bars indicate ± SD, n = 3. Statistical comparisons are by Student’s t-test: **p<0.01 (F) Schematic showing the approach to recruit clients into condensates in yeast cells. SUMO_10_-SIM_6_ was expressed in cells to form condensates. ERK1 and ELK1 (clients) were tagged with SIM for recruitment into condensates. ERK1 and ELK1 were also fused with GFP and mCherry respectively. (G) Recruitment to condensates increases phosphorylation levels *in vivo*. Quantification of ELK1^DM^ phosphorylation in yeast cells. Top panel, micrographs of control *S. cerevisiae* cells with various combinations of ERK1^DM^ (green), ELK1 (red) and scaffold controls or SUMO_10_-SIM_6_ condensates (no fluorescent tag). SIM tags enabled client recruitment into SUMO_10_-SIM_6_ condensates. Scale bar = 5 μm. Below, representative western blots for (from top to bottom) the ELK1-S383 phosphoepitope, total ELK1, the activation-loop phosphates on the ERK1 kinase (pT202/pY204), and tubulin loading control. Red arrowheads indicate hypershifted bands. Quantification of ELK1 S383 phosphorylation (pS383) and the hyperphosphorylation (Hypershift / Tubulin) are shown in bottom graphs. For S383 phosphorylation, band intensities of total phosphorylated ELK1 were normalized to total ELK1 levels, and this value was further normalized to the phosphorylation level of ELK1 in the control strain (leftmost) expressing only ELK1. For hyperphosphorylation (Hypershift / Tubulin) quantification, the intensity signal of hypershifted band (detected by anti-p-ELK1-S383) was normalized to tubulin levels in the bottom graph, and the values were further normalized to the leftmost strain expressing ELK1 only. Error bars indicate ± SD, n = 3, Statistical comparisons are by Student’s t-test: *p<0.05, N.S., not significant.

The first clients we tested were the human protein kinase ERK1 and one of its substrates, the C-terminus of ELK1. We took advantage of an R84S mutant of ERK1 that does not require activation by upstream activating kinases (i.e. MEK) and is constitutively active both *in vitro* and when expressed *in vivo* (Figure 1B) (Levin-Salomon et al., 2008). The ELK1 C-terminus contains multiple ERK1 consensus phosphorylation motifs (Mylona et al., 2016) and two docking motifs (D site and F site) that bind to the surface of ERK1. We tested the wild-type (WT) ELK1 C-terminus (Figure 1C, residues 206-428, hereafter referred to as ELK1) and a version in which these two docking motifs were mutated (hereafter referred to as ELK1^DM^). To enable rapid recruitment of clients into condensates, we fused an FKBP12 domain to the N-terminus of SIM_6_, and an FRB domain and fluorescent protein (for visualization) to the N-terminus of each client. When FKBP12 is bound to the small macrolide rapamycin, it subsequently binds to FRB with high-affinity. Therefore, as expected, addition of rapamycin recruited FRB-GFP-ERK1 and FRB-mCherry-ELK1into the condensates of SUMO_10_ + FKBP12-SIM_6_ (Figure 1A & 1D, Figure 1 Supplement 1A). We simultaneously measured the fluorescence intensity of GFP-ERK1 and mCherry-ELK1 inside and outside condensates to calculate the partition coefficient of each client. Rapamycin addition increased the partition coefficient of GFP-ELK1 within condensates (Figure 1D).

We compared the rate of phosphorylation of ELK1 after recruitment into condensates (using rapamycin) to the rate without (using DMSO as a solvent control). We measured the initial rate at which Erk1 catalyzed the incorporation of radiolabel from γ-^32^P-ATP into ELK1 during 5 minutes. Quantification showed that recruitment into condensates by rapamycin addition increased rates or phosphorylation of ELK1^WT^ by 22% compared to control (Figure 1E, Figure 1 Supplement 1C), suggesting that recruitment of ERK1 and ELK1 into SUMO_10_ + SIM_6_ condensates accelerated the reaction. The ELK1^DM^ mutant had lower baseline phosphorylation, as expected, but the relative increase in phosphorylation upon recruitment into SUMO_10_ + SIM_6_ condensates was greater than ELK1^WT^, nearly doubling (Figure 1E, Figure 1 Supplement 1C). Together these results indicate that recruitment of ERK1 kinase and the ELK1 substrate into a condensed phase accelerates the rate of phosphorylation, especially for substrates that lack docking motifs. The reaction acceleration in this experiment appears to be relatively small (22% for ELK1^WT^, 91% for ELK1^DM^), however, the condensate phase only constituted a tiny fraction of the total reaction volume (less than 3%). Therefore, as previously reported for similar experiments (Peeples and Rosen, 2021), we believe the actual acceleration in condensates is far greater than apparent in the bulk reaction.

### Recruitment to synthetic condensates increases phosphorylation *in vivo*

We next investigated whether synthetic condensates could potentiate phosphorylation *in vivo*. We used the budding yeast *Saccharomyces cerevisiae* as our model system. In contrast to our *in vitro* system, which was a coacervate of independent SUMO_10_ and SIM_6_ polypeptides, we expressed a single polypeptide consisting of SUMO_10_ fused to SIM_6_ (SUMO_10_-SIM_6_) (Banani et al., 2016). This approach ensures that there are always excess SUMO binding sites available within the SUMO_10_-SIM_6_ condensate. Rapamycin causes important changes to the biophysical properties of *S. cerevisiae* (Delarue et al., 2018, and see figure 6 below), therefore, we did not use the FKBP12/FRB system to recruit clients to these condensates *in vivo*. Instead, we tagged the constitutively active ERK1^R84S^ kinase and its substrate ELK1 at their C-termini with a single SIM motif. The SUMO_10_-SIM_6_ condensates have four excess SUMO binding sites per scaffold, enabling recruitment of these SIM-tagged clients (Figure 1F). We expressed combinations of SIM-tagged WT ELK1 or ELK1^DM^ with SUMO_10_-SIM_6_ in *S. cerevisiae* and analyzed phosphorylation levels by western blotting using an ELK1 Serine-383-phospho-specific antibody. There was no increase in phosphorylation levels in the absence of SIM-tagged ERK1. To control for scaffolding effects we also used a SUMO dimer (SUMO_2_) that did not form condensates. We observed some basal phosphorylation of ELK1 even without ERK1 expression, indicating cross-talk between orthogonal yeast and human proteins. Both ELK1^WT^ and ELK1^DM^ phosphorylationat S383 was increased when co-recruited with ERK1 into condensates (3.8-fold and 4.9-fold respectively, p < 0.05, Figure 1G top graph, Figure 1 Supplement 2). Again, ELK1^DM^ showed a greater relative increase in phosphorylation upon condensate recruitment. As expected, overall phosphorylation of ELK1^WT^ by ERK1 outside of condensates was higher than for ELK1^DM^, though recruitment to condensates still increased phosphorylation (Figure 1 supplement 2). SUMO_2_ expression did not significantly increase ELK1^WT^ or ELK1^DM^ phosphorylation, indicating that recruitment to SUMO_10_-SIM_6_ condensates and not simply scaffolding effects (i.e. the simultaneous binding of ERK1 and ELK1 to adjacent SUMO domains in the dispersed solute phase) were required for increased phosphorylation. Due to the nature of the western blotting procedure, these phosphorylation levels are population averages and represent pseudo-steady-state values. Therefore, we cannot determine precise rates. However, taken together, and combined with the *in vitro* results above, these controls and experiments are most consistent with the simple interpretation that phosphorylation rates are increasing in condensates *in vivo*.

### Recruitment to synthetic condensates leads to novel multi-site phosphorylation *in vivo*

In addition to increased intensity of phospho-ELK1 bands we also noticed prominent slower migrating species specifically in conditions of condensate recruitment. ELK1 is known to be phosphorylated at multiple sites by ERK1. Therefore, we hypothesized that the slowly migrating bands on the western blot were due to multi-site phosphorylation. To test that the hypershifted bands were due to phosphorylation, we treated cell lysates with lambda phosphatase. Lambda phosphatase treatment resulted in almost complete loss of the hypershifted bands, while phosphatase inhibitor addition maintained the hypershifted band, suggesting that phosphorylation caused the slow migration pattern (Figure 1 Supplement 3A). ELK1 contains eleven serines or threonines immediately followed by proline, corresponding to consensus ERK1 phosphorylation sites (Cruzalegui et al., 1999). Substitution all of these serines/threonines (Figure 1 Supplement 3B) with alanine (ELK1-11A mutant) led to loss of almost all hypershifted bands, suggesting that phosphorylation at these sites was responsible for the slow migration pattern (Figure 1 Supplement 3D). Quantification of slow-migrating bands indicated a 220-fold increase in hyperphosphorylation when ELK1^DM^ was recruited to condensates (Figure 1G, bottom graph, p < 0.05) and a 380-fold increase in the hypershifted band for ELK1^WT^ (Figure 1, supplement 2, bottom graph, p < 0.05). ELK1 was previously shown to be phosphorylated at different sites with different kinetics (Mylona et al., 2016). Serine 383 (Serine 384 in mouse Elk-1) is the most rapidly phosphorylated residue, perhaps leading to partial reaction saturation even when ELK1^WT^ and ERK1 are co-expressed without SUMO_10_-SIM_6_, and explaining the modest increase in phosphorylation of this residue when ELK1^WT^ is recruited to condensates. On the other hand, condensate recruitment seems to have a far more pronounced effect on the phosphorylation of residues that are normally inefficiently phosphorylated (Mylona et al., 2016).

### Condensates facilitate new links between kinases and substrates

Our results so far suggested that, when recruited to synthetic condensates, ERK1 can efficiently phosphorylate ELK1 without docking motifs and on suboptimal phosphoacceptor residues (Figures 1E and 1G, Figure 1 Supplement 3.) We therefore hypothesized that recruitment into condensates might create a permissive environment for phosphorylation, potentially expanding kinase specificity. To test this idea, we investigated whether other kinases that did not evolve with ELK1 as a substrate could phosphorylate ELK1 when recruited to SUMO_10_-SIM_6_ condensates. We first selected Fus3p (hereafter Fus3), a yeast mitogen activated protein kinase (MAPK), which is the yeast kinase most closely evolutionarily related to ERK1. We tagged the endogenous *FUS3* gene with a double SIM tag (gene product hereafter, Fus3-2xSIM) to recruit it into SUMO_10_-SIM_6_ condensates (Figure 2A). Western blot results showed that ELK1-SIM phosphorylation on S383 increased ∼6-fold when Fus3-2xSIM was co-localized in SUMO_10_-SIM_6_ condensates (Figure 2B, top graph, p < 0.01). This result indicates that basal activity of Fus3-2xSIM can drive significant phosphorylation of a novel substrate in condensates. The α factor mating peptide triggers a signal transduction cascade that results in increased Fus3 activity (Figure 2A) (Bardwell, 2004). The level of ELK1-SIM S383 phosphorylation was further increased ∼12-fold above background upon activation of Fus3 by addition of α factor for 60 min (Figure 2B, top graph, p < 0.01). Furthermore, hypershifted bands were even more strongly induced by α factor (∼250-fold, p < 0.05, Figure 2B, bottom graph). Expression of ELK1-SIM and Fus3-2xSIM together with a simple SUMO_2_ scaffold did not result in increased phosphorylation of ELK1, regardless of α factor addition. These results indicate that a novel kinase-substrate connection can be induced by recruitment to a condensate, and that information from upstream signaling can still be received within this condensed phase.

**Figure 2:**
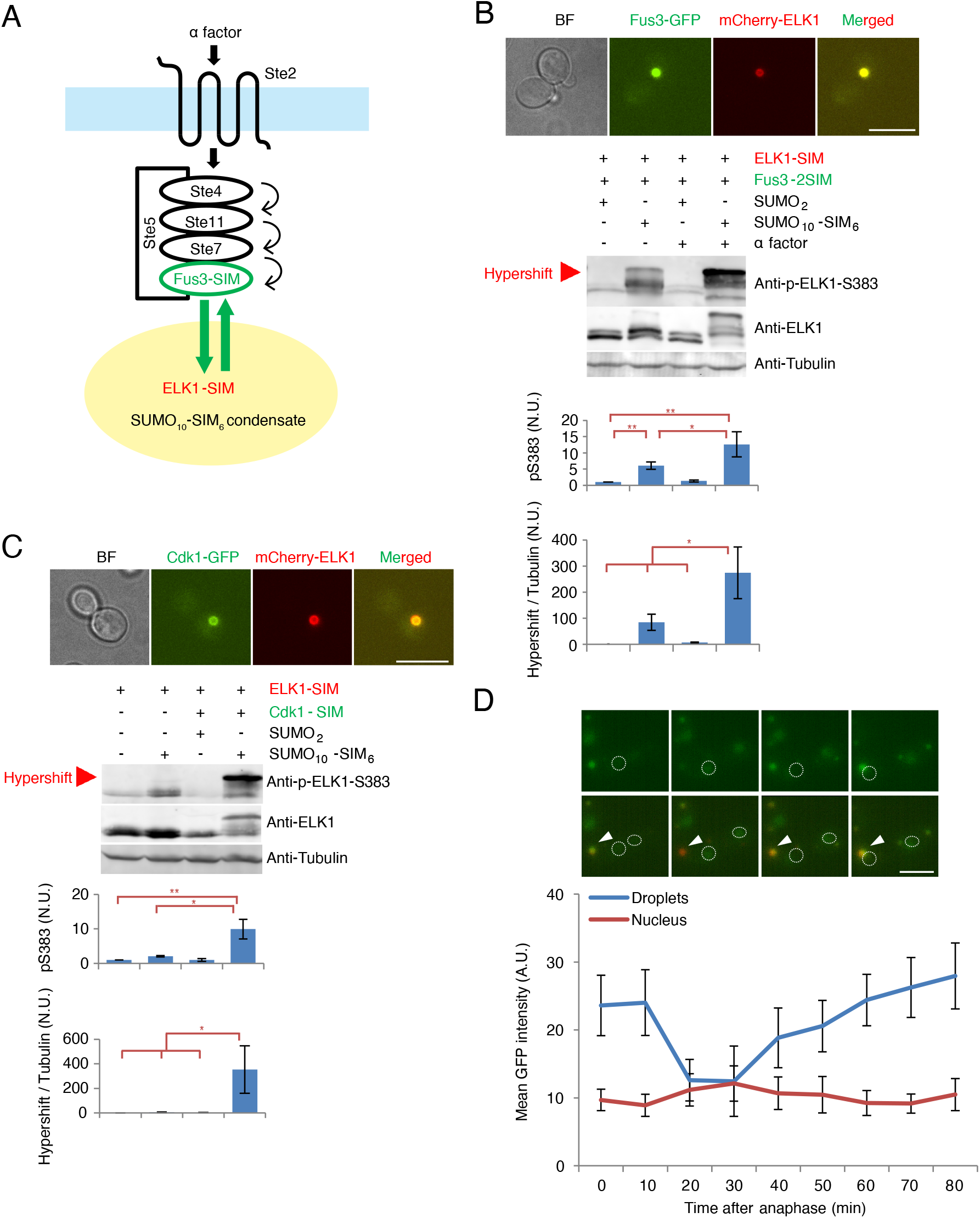
Condensates facilitate new links between kinases and substrates. (A) Schematic illustrating the approach to test ELK1 phosphorylation by Fus3 in condensates, and the pathway for Fus3 activation by α factor in yeast. Fus3 was recruited to SUMO_10_-SIM_6_ condensates with a 2xSIM tag. (B - C) Co-localization of Fus3 (B) or Cdk1 (C) with ELK1 in SUMO_10_-SIM_6_ condensates leads to increased phosphorylation of ELK1, scale bar = 5 μm. Cells were treated with 2 µM alpha factor for 1h to activate Fus3 (B).In the middle panels of (B) and (C), representative western blots are shown for (from top to bottom) the ELK1-S383 phosphoepitope, total ELK1, and tubulin loading control. Red arrowheads indicated hypershifted bands. Quantification of ELK1 S383 phosphorylation (pS383) and the hyperphosphorylation (Hypershift / Tubulin) are shown in bottom graphs. For S383 phosphorylation, band intensities of total phosphorylated ELK1 were normalized to total ELK1 levels. For hyperphosphorylation (Hypershift / Tubulin) quantification, the intensity signal of hypershifted band (detected by anti-p-ELK1-S383) was normalized to tubulin levels in the bottom graph. The value was further normalized to the phosphorylation level or hyperphosphorylation level of Elk1 in the control strain (left) expressing ELK1, Fus3 and SUMO_2_ (B) or ELK1 only (C). Error bars indicate ± SD, n = 3. Statistical comparisons are by Student’s t-test *p<0.05, **p<0.01. (D) The NLS-GFP-2xWW dynamically changes localization during mitosis. Log-phase cells were immobilized on a 384-well imaging plate, and z-stacks were acquired every 10 minutes. Average projections of the GFP channel (GFP-2xWW reporter, top) and a merge of the mCherry and GFP channels (GFP-2xWW, green; mCherry-ELK1, red) are shown. The white dotted line indicates the position of the nucleus as detected by a 2xNLS-BFP reporter. White arrowheads indicate the reporter in condensates, scale bar = 5 μm. The mean intensity of GFP in condensates and in the nucleus were quantified and are plotted as a line graph, n = 10 cells, error bars indicate ± SD.

Next we tested if a more evolutionarily distant kinase could phosphorylate ELK1 within condensates. We chose the Cyclin Dependent Kinase Cdk1 (encoded by the *CDC28* gene in *S. cerevisiae*). Cdk1 is a member of the CMGC super-family of kinases and is still proline-directed, but is structurally and functionally distinct from MAP kinases, and recognizes different docking motifs (Howard et al., 2014; Mok et al., 2010; Schulman et al., 1998). We tagged the endogenous Cdk1 with a single SIM peptide (Cdk1-SIM) and observed robust recruitment into SUMO_10_-SIM_6_ condensates (Figure 2C, top). Co-recruitment of ELK1-SIM and Cdk1-SIM led to high levels of phosphorylation of ELK1, both in terms of S383 phosphorylation (∼10-fold increase, p < 0.01, Figure 2C, top graph) and levels of hypershifted bands (∼300-fold increase, p < 0.05, Figure 2C, bottom graph). Expression of ELK1-SIM and Cdk1-SIM together with a simple SUMO_2_ scaffold did not increase ELK1 phosphorylation over background levels. Cdk1 is the master regulator of the cell division cycle (Malumbres, 2014). Cdk1 activity is highest during mitosis and is then inactivated during mitotic exit by degradation of cyclins (Enserink and Kolodner, 2010). In addition, the CDC14 phosphatase that counteracts Cdk1 is specifically activated during mitotic exit (Stegmeier and Amon, 2004). Therefore, substrates of Cdk1 are dynamically phosphorylated and dephosphorylated as the cell cycle progresses. To further investigate the ability of synthetic condensates to create novel kinase connections while allowing dynamic regulation, we built a reporter system to visualize phosphorylation of ELK1 by Cdk1 in real-time in single cells.

To create a live-cell reporter of ELK1 phosphorylation, we fused two WW domains from the *Homo sapiens* PIN1 protein (Peptidyl-prolyl cis-trans isomerase NIMA-interacting 1) to GFP (GFP-2xWW). The WW domain specifically interacts with phosphorylated serine-proline motifs (Verdecia et al., 2000). We hypothesized that ELK1 phosphorylation within synthetic condensates would lead to strong interactions with this reporter, thus leading to its recruitment to the cytoplasmic condensates (Figure 2 Supplement Figure 1A). Indeed, we found that coexpression of ERK1-SIM and ELK1-SIM led to the translocation of the GFP-2xWW reporter into SUMO_10_-SIM_6_ condensates (Figure 2 Supplement 1B, top). As a control, we expressed a catalytically dead ERK1 mutant (ERK1-K71R) and found very little recruitment of the reporter into SUMO_10_-SIM_6_ (Figure 2 Supplement 1B, bottom). These results suggested that the reporter was recruited to synthetic condensates in a phosphorylation-dependent manner.

We next modified the reporter to attempt to reveal dynamic phosphorylation of ELK1 by Cdk1. We added an SV40 nuclear localization signal (NLS) to the N-terminus (NLS-2xWW-GFP), such that, when the reporter and ELK1 were coexpressed with SUMO_10_-SIM_6_ but Cdk1 was not tagged with SIM, most of the reporter remained nuclear, and very little was recruited to condensates. However, when Cdk1 was tagged with SIM, a significant number of cells had strong reporter recruitment to condensates (Figure 2 Supplement 2), suggesting that ELK1 was phosphorylated by Cdk1 within condensates. We next examined the dynamics of reporter recruitment to condensates during the cell cycle by time lapse imaging (Figure 2D). Reporter intensity within condensates was highest immediately prior to anaphase, the point at which Cdk1 activity is highest (Enserink and Kolodner, 2010). Approximately 20-30 minutes after anaphase, the reporter intensity was dramatically reduced within condensates, and relocalized to the nucleus (Figure 2D, supplemental movie 1). This corresponds to the timing of mitotic exit, when Cdk1 is rapidly inactivated by cyclin degradation and expression of the Cdk1 inhibitor Sic1 (Enserink and Kolodner, 2010). In addition, the phosphatase Cdc14 is activated at this time, leading to rapid dephosphorylation of phosphorylated S-P and T-P motifs (Visintin et al., 1998). Subsequently, after 10-20 minutes, a time when Cdk1 activity is again rising, the reporter intensity in condensates also started to rise (Figure 2D, Figure 2 Supplement 3). Collectively, these results suggest that recruitment of Cdk1-SIM to a synthetic condensate can create a new kinase-substrate connection to ELK1, and that phosphorylation within SUMO_10_-SIM_6_ condensates can change on the minute time-scale, with dynamics that closely follow the changes in Cdk1 activity during the cell cycle.

We next wondered if recruitment to condensates would relax the primary specificity of the kinase, enabling phosphorylation of a broader range of peptides. First, we used a short peptide flanking serine383 in ELK1 as a substrate (residue 370-394, referred to as min383 hereafter), to completely remove the D box and F box docking sites. We fused this peptide and ERK1 to a 5xSIM tag to obtain strong enrichment within condensates (Figure 2 Supplement 4A). As shown in Figure 2 Supplement 4B, 4C and 4D, serine383 of the min383 ELK1 substrate was highly phosphorylated when clients were co-localized in SUMO_10_-SIM_6_ condensates, compared to the low phosphorylation in the strains without condensates, suggesting that docking motifs are not required for ERK1 phosphorylation within condensates. Interestingly, condensates were also formed when 5xSIM-tagged-clients were co-expressed with scaffold SUMO_2_ (Figure 2 Supplement 4C), resulting in increased min383 phosphorylation, and these condensates also drove phosphorylation.

We noticed that there were multiple hypershifted bands, accounting for almost 50% of total min383 (Figure 2 Supplement 4D) within SUMO_10_-SIM_6_ condensates. These bands disappeared after lambda phosphatase treatment (Figure 2 Supplement 5A), suggesting that phosphorylation caused this slow migration pattern. There is only one additional ERK1 consensus phosphoacceptor site (S/T-P motif, serine389) in min383 (Figure 2 Supplement 5B), therefore we hypothesized that other non-consensus sites could also be phosphorylated. We generated several point mutant and truncation constructs to identify the phosphorylated residues (Figure 2 Supplement 5C). Western blot results showed that the slowly migrating bands almost completely disappeared in a truncation construct that deleted several serines at the N- and C-terminus. Deletion of four consecutive serines at the N-terminus almost completely abolished slowly migrating bands (Figure 2 Supplement 5D), suggesting that these four non-consensus serines were phosphorylated by ERK1. Thus, ERK1 can phosphorylate non-consensus. Sequences within condensates.

Extending from these observations, we next investigated the ability of Cdk1 to phosphorylate non-canonical substrates within condensates. Cdk1 is also a proline-directed kinase, so we generated a construct that fused short peptides from the human P53 and RPS6 proteins (Figure 2 Supplement 6A and 6B, referred to PR hereafter), none of which contain any known primary specificity determinants for Cdk1 (Mok et al., 2010). There are also commercial phospho-specific antibodies available to detect phosphorylation of these sites. Western blot results showed that peptides from serine9 and serine37 in P53 were more highly phosphorylated by Cdk1 in condensates compared to without condensates (Figure 2 Supplement 6C and 6D). Serine235/236 in RPS6 was also phosphorylated by Cdk1 in condensates. Collectively, these results show that the primary specificity of ERK1 and Cdk1 is expanded when recruited to condensates.

## Multiple factors contribute to hyperphosphorylation in condensates

A major advantage of synthetic biology is the possibility of systematically varying parameters in an attempt to understand general principles. Therefore, we varied the properties of both the. condensate scaffold proteins, and the clients to investigate which factors impact phosphorylation of ELK1 by ERK1 within condensates. We focused on hyperphosphorylation, which is more dynamically affected by recruitment to condensates than phosphorylation at S383 alone.

First, we characterized the effect of client binding affinity on phosphorylation in condensates. A single substitution in the SIM peptide (substitution of Isoleucine 9 with proline, referred to as SIM^I9P^ hereafter) decreases the affinity of the interaction between SIM and SUMO (Namanja et al., 2012). We tagged the ELK1 substrate with either WT SIM (SIM^WT^) or SIM^I9P^, while the ERK1 kinase was always tagged with SIM^WT^. The concentration of ELK1-SIM^I9P^-tagged substrate was dramatically reduced in SUMO_10_-SIM_6_ condensates (Figure 3B and Figure 3 Supplement 1). Western blotting showed that hyperphosphorylation of ELK1-SIM^I9P^ in condensates was reduced by almost 80% compared to ELK1-SIM^WT^ (Figure 3B and Figure 3 Supplement 2). We hypothesized that the reduced substrate concentration led to the decreased hyperphosphorylation. To test this idea, we used a stronger promoter to increase the expression level of ELK1-SIM^I9P^ and restore the concentration of ELK1-SIM^I9P^ in condensates. Indeed, this higher substrate concentration within condensates mostly restored ELK1-SIM^I9P^ hyperphosphorylation. Collectively, these results suggested that client affinity affects hyperphosphorylation in condensates, at least partly through client concentration.

**Figure 3:**
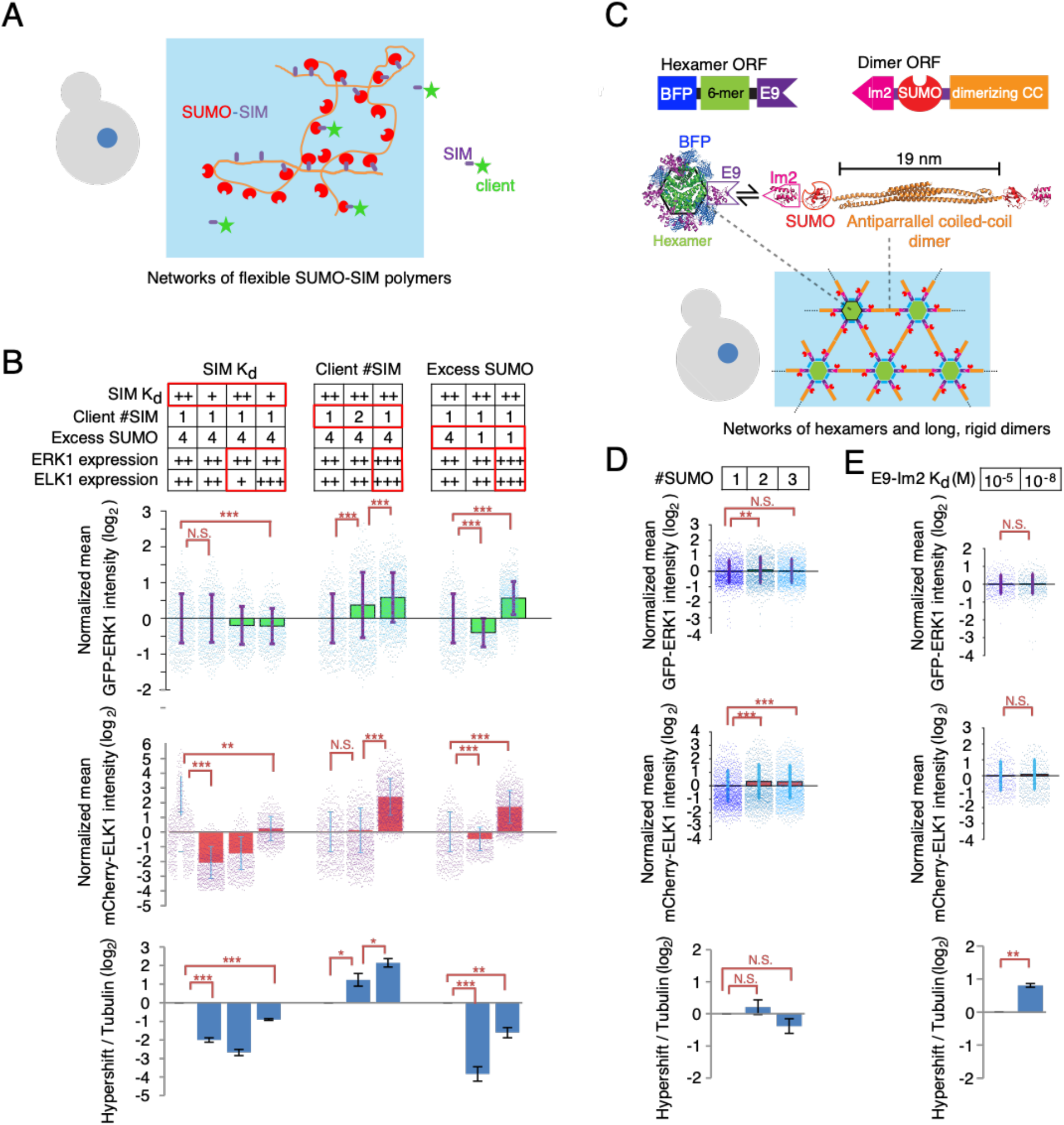
Multiple factors contribute to hyperphosphorylation in condensates. (A) Schematic of SUMO_n_-SIM_n_ condensates in *S. cerevisiae* cells. (B) We varied the interaction strength between clients and condensates using variants of the SIM motif with low K_d_ (higher affinity, ++) or high K_d_ (lower affinity, +). We varied the number of SIM tags on the client proteins (either one or two SIMs), and the availability of excess client binding sites on scaffolds by using either SUMO_10_-SIM_6_ (4 excess SUMO per scaffold) and SUMO_7_-SIM_6_ (1 excess SUMO per scaffold). We also varied the promoters for kinase and substrate at obtain high (+++), medium (++), or low expression levels (+) to modulate client concentration within condensates. The relative concentrations of GFP-ERK1-SIM and mCherry-ELK1-SIM within condensates were estimated based on fluorescence intensities from average projections of z-stacks, and are shown on a log_2_ scale in the red and green bar graphs, where each point is the quantification of a condensate. Bottom (blue bar graph) shows quantification of hyperphosphorylation levels from western blots. Client concentration and ELK1 hyperphosphorylation values were normalized to the median of the leftmost strain expressing medium amounts of clients tagged with a single wild-type SIM and co-expressing a SUMO_10_-SIM_6_ condensate. (C) Schematic showing the design of a synthetic two-component condensate system for expression in *S. cerevisiae* cells. One component consists of a homodimerizing scaffold fused to a SUMO domain (or multiple SUMO domains) that can recruit SIM-tagged client proteins, and the Im2 domain. The other component consists of a homohexamerizing scaffold fused to the blue fluorescent protein (BFP) and the E9 domain. The Im2 and E9 domains interact with one another. The dimer-forming component is too long (> 18 nm) and rigid to bind onto the same hexameric assembly more than once. Therefore, an extended network of interactions occurs, leading to the formation of a relatively rigid condensate with SUMO domains constrained apart from one another. (D-E) Characterization of the effect of the number of SUMO domains on each dimer-forming subunit, and scaffold dynamics within the condensate. (D) One, two, or three SUMO domains were inserted into the dimer-forming component. Client concentrations within the condensate were estimated as above and normalized to the concentration within condensates containing a single SUMO on each dimer-forming subunit. Bottom (blue bar graph) shows quantification of hyperphosphorylation levels from western blots. (E) Characterization of the effect of the affinity of the Im2/E9 interaction on hyperphosphorylation within synthetic two component condensates. Two Im2 variants were used, the wild-type domain nanomolar affinity (K_d_ ∼ 1.2 × 10^−8^ M), while a weaker variant had an affinity in the micromolar (K_d_ ∼ 3 × 10^−5^ M) range. Client concentrations within condensates were estimated as above and normalized to the concentration within condensates with a micromolar Im2/E9 interaction. Bottom (blue bar graph) shows quantification of hyperphosphorylation levels from western blots. In all western blots, the intensities of hypershifted ELK1 bands were normalized to tubulin expression. n = 3. The hyperphosphorylation level of ELK1 in the leftmost strain was set as 1. Error bars indicate ± SD; statistics by Student’s t-test: *p < 0.05; **p < 0.01; ***p < 0.001; N.S., not significant.

When ELK1-SIM^I9P^ was present at equivalent concentration to ELK1-SIM^WT^, there was still a two-fold decrease in hyperphosphorylation. One possible explanation is that the binding kinetics of client to the condensate SUMO domains plays an important role. The unbinding rate (k_off_) of ELK1-SIM^I9P^ is likely to be higher than ELK1-SIM^WT^. An alternative way to modulate k_off_ is to increase the valency of SIM motifs on the clients. We therefore fused both kinase and substrate with two SIM-tags (2xSIM) to investigate the effect of client valence on hyperphosphorylation in condensates. The 2xSIM-tag led to a slightly higher (∼ 20%) ERK1-GFP concentration in condensates compared the single SIM-tag (Figure 3B and Figure 3 Supplement 1). However, western blot results show that the hyperphosphorylation level of 2xSIM-tagged clients was almost doubled (Figure 3B and Figure 3 Supplement 2). We compared the 2xSIM-tagged substrate phosphorylation to strongly overexpressed of both kinase and substrate with a single SIM-tag. This strong overexpression led to higher client concentration in condensates and slightly higher ELK1 hyperphosphorylation. Therefore, binding kinetics of clients modulates the degree of hyperphosphorylation within condensates.

We next investigated how the properties of the synthetic condensates could impact hyperphosphorylation. The presence of excess SUMO domains within the SUMO_10_-SIM_6_ condensates has been shown to be important for client recruitment (Banani et al., 2016). We therefore tested the effect of reducing the number of free SUMO domains by comparing SUMO_7_-SIM_6_ to SUMO_10_-SIM_6_ condensates. Client recruitment into SUMO_7_-SIM_6_ condensates was only slightly lower than into SUMO_10_-SIM_6_ condensates (Figure 3B and Figure 3 Supplement 1). However, ELK1 hyperphosphorylation within SUMO_7_-SIM_6_ condensates was reduced by almost 90% (Figure 3B and Figure 3 Supplement 2). Next, we asked whether we could restore hyperphosphorylation in SUMO_7_-SIM_6_ condensates by increasing client concentration in condensates. Interestingly, hyperphosphorylation was still ∼3-fold lower in strains expressing SUMO_7_-SIM_6_ condensates when the expression of both clients was increased such that client concentration in SUMO_7_-SIM_6_ was higher than in SUMO_10_-SIM_6_ condensates. This suggests that the excess SUMO domains in condensates potentiates hyperphosphorylation through a mechanism beyond simple mass-action.

Taken together, these results suggest that client concentration, binding kinetics, and the availability of excess SUMO domains in the condensate are all important determinants of the efficiency of phosphorylation within condensates. Interestingly, it seems that the properties of the condensate *per se* are the most important factor, in particular the presence of a large excess of client binding sites.

### Condensate scaffold flexibility is crucial for hyperphosphorylation

The SUMO-SIM condensates are based on highly flexible polymers that can assemble in many conformations. We wondered whether this flexibility, which appears to be a frequent feature of biological condensates (Alberti and Hyman, 2021; Hastings and Boeynaems, 2021; Li et al., 2012), was important for the potentiation of phosphorylation. We took advantage of a recently described design strategy (Heidenreich et al., 2020) based on a well-structured synthetic two component system (Figure 3C). One component consists of the protein O27018 from *Methanobacterium thermoautotrophicum*, which is a self-assembling homo-hexamer of unknown function (PDB 3BEY). The other component is a homo-dimer consisting of a long (18 nm), rigid, anti-parallel coiled-coil domain from TRIM25 (PDB 4LTB). Interaction between these two components was engineered using a well-characterized heterodimeric interaction between a mutated endonuclease domain of the bacterial toxin colicin E9 (henceforth simply E9), and the immunity protein, Im2. We fused E9 to the C-terminus of the hexamer and Im2 to the N-terminus of the dimer. An important design feature is that the dimer is too long to bind twice to the same hexamer, thereby making lattice assembly within the condensate more predictable. We additionally fused a SUMO domain to the N-terminus of the dimer. The topology of the dimer is such that the N-termini are distant from one another at the opposite ends of each dimer. In summary, this synthetic condensate is predicted to assemble in a limited set of local geometries from rigid components that hold individual SUMO domains relatively distant from one another.

When expressed in yeast, the synthetic two-component formed condensates that recruited ELK1-SIM and ERK1-SIM clients (Figure 3 Supplement 3). We found that recruitment of ELK1-SIM and ERK1-SIM to SUMO domains within this ordered condensate led to a slight increase in total phosphorylation levels at S383 (∼50%) and only mild increase hyperphosphorylation (∼35 fold) (Figure 3 Supplement 4A and 4B), less than in SUMO-SIM condensates. Thus, scaffold flexibility may be an important feature that allows efficient phosphorylation within condensates.

### The presence of up to three adjacent client binding sites is insufficient to drive efficient phosphorylation in ordered condensates

A major feature of the two-component system is that the SUMO binding sites are held at a slight distance from one another within a relatively rigid network. This is distinct from the SUMO-SIM condensates, where multiple adjacent SUMO domains can be available for client binding in a single polymer, and the scaffold flexibility potentially allows for even more extensive clusters of SUMOs to occur. Indeed, the fact that the availability of excess SUMO domains has the largest effect on the efficiency of hyperphosphorylation (Figure 3B, right) suggests that the availability of local densities of client binding sites is crucial for condensed-phase signaling. Control experiments demonstrate that dimers of SUMO in solution are insufficient for reaction acceleration (Figure 1G, scaffolding control, Figure 1 Supplement 2, Figures 2B and 2C, scaffolding control.) To gain further insights, we next investigated if adding pairs or triads of SUMO domains to the dimer in the two-component condensate could better drive condensed-phase phosphorylation. We screened for colonies with similar client concentrations within condensates to simplify interpretation. We fused the dimer component to 2xSUMO and 3xSUMO. Again, each 2xSUMO or 3xSUMO is held 18 nm distant from its partner by the rigid antiparallel coiled-coil of the dimer, and when bound to hexamer can be within (9-12 nm) of a set of SUMO sites on an adjacent dimer head (Figure 3C). We found no further increase in S383 phosphorylation or hyperphosphorylation of ELK1 with 2xSUMO or 3xSUMO relative to 1xSUMO within these rigid condensates (Figure 3D). These results suggest that, even when a huge concentration of 3xSUMO domains is present within a condensate, the inability of the rigid scaffold network to generate larger SUMO clusters may limit the efficiency of this condensate as a catalyst of hyperphosphorylation. Therefore, we conclude that local pockets of high binding site concentration (>3 SUMO sites), and perhaps scaffold polymer dynamics *per se*, play an important role in condensed phase signaling.

### Scaffold material properties do not greatly affect phosphorylation within ordered condensates

A useful feature of the two-component system is the availability Im2 variants with varying affinity to E9 (Heidenreich et al., 2020; Li et al., 1998). We took advantage of these mutations to create condensates with two interaction strengths between the two scaffold components, either in the micromolar (K_d_ ∼ 3 × 10^−5^ M) or nanomolar (K_d_ ∼ 1.2 × 10^−8^ M) range. Im2 variants with a range of affinity towards E9 have been shown to create liquid and solid-like condensates respectively (Heidenreich et al., 2020). We screened for yeast strains expressing each of these variants that had similar condensate sizes and concentration of clients in the condensates (Figure 3E; Figure 3 Supplement 4C). Western blotting showed a slight increase (∼2-fold) in the strain expressing the high affinity (10^−8^M) Im2 variant over that of the lower affinity (10^−5^M) variant (Figure 3E, Figure 3 Supplement 4D). This result suggests the material states and dynamics of the condensed scaffold only play a minor role in condensed-phase phosphorylation within these well-structured networks.

### Synthetic condensed phase signaling can respond to osmotic compression

Osmotic compression has multiple effects on cell, including reduced cell size, increased macromolecular crowding, and decreased molecular diffusion (Joyner et al., 2016; Miermont et al., 2013). We have previously shown that osmotic compression can influence condensate formation through macromolecular crowding effects (Delarue et al., 2018). On a microscopic scale, the assembly, dynamics, and network structure of condensates could all be impacted by macromolecular crowding, as could the dynamics of client interactions and motion within the condensed phase. Client molecules exist in two states within condensates. Some client molecules are in the fluid phase that permeates the condensate and not engaged with binding sites, while others are bound to the condensate scaffolds through SIM interactions between the SIM on the clients and SUMO on the scaffold. As macromolecular crowding impacts the condensate network, both the motion through the fluid phase and the binding kinetics of clients could be impacted. Thus, it is possible the phosphorylation within condensates could be sensitive to changes in the biophysical properties of the cell and provide a mechanism to convert physical signals such as macromolecular crowding to chemical signals. We took advantage of our synthetic condensates to test this idea and investigate factors important for this physical sensing.

We used *hog1Δ* strains that are deleted for the main osmotic stress response kinase. This mutation prevents adaption to osmotic pressure, and avoids the issue of changes in signaling due to *Hog1* activation upon osmotic stress. We tested condensates that varied in the two factors that we previously determined to have the greatest effect on phosphorylation efficiency: condensate network flexibility (the relatively rigid two-component condensate versus flexible SUMO-SIM condensates), and the availability of excess SUMO (i.e. SUMO_10_-SIM_6_ versus SUMO_7_-SIM_6_). The clients were the kinase GFP-ERK1-SIM and the substrate mCherry-ELK1-SIM.

First, we tested the effect of osmotic compression of SUMO_10_-SIM_6_ condensates. We osmotically compressed cells with 1M sorbitol for one hour and quantified client recruitment, condensate size, and hyperphosphorylation of ELK1. The average condensate area decreased by around 35%, likely due to oncotic compression, and recruitment of GFP-ERK1-SIM increased ∼70% (Figure 4 supplement 1), but ELK1 hyperphosphorylation was only increased about 10% (Figure 4 supplement 2). We reasoned that this could be because SUMO_10_-SIM_6_ condensates already give the most efficient hyperphosphorylation of any synthetic condensate. We therefore next tested the effect of osmotic compression of SUMO_7_-SIM_6_ condensates. The change in condensate area and client concentration was similar to that in SUMO_10_-SIM_6_ condensates (35% area decrease and ∼70% increase in GRP-ERK1-SIM, Figure 4 Supplement 1). However, the degree of ELK1 hyperphosphorylation in SUMO_7_-SIM_6_ condensates was increased ∼2-fold (Figure 4 supplement 2).

**Figure 4:**
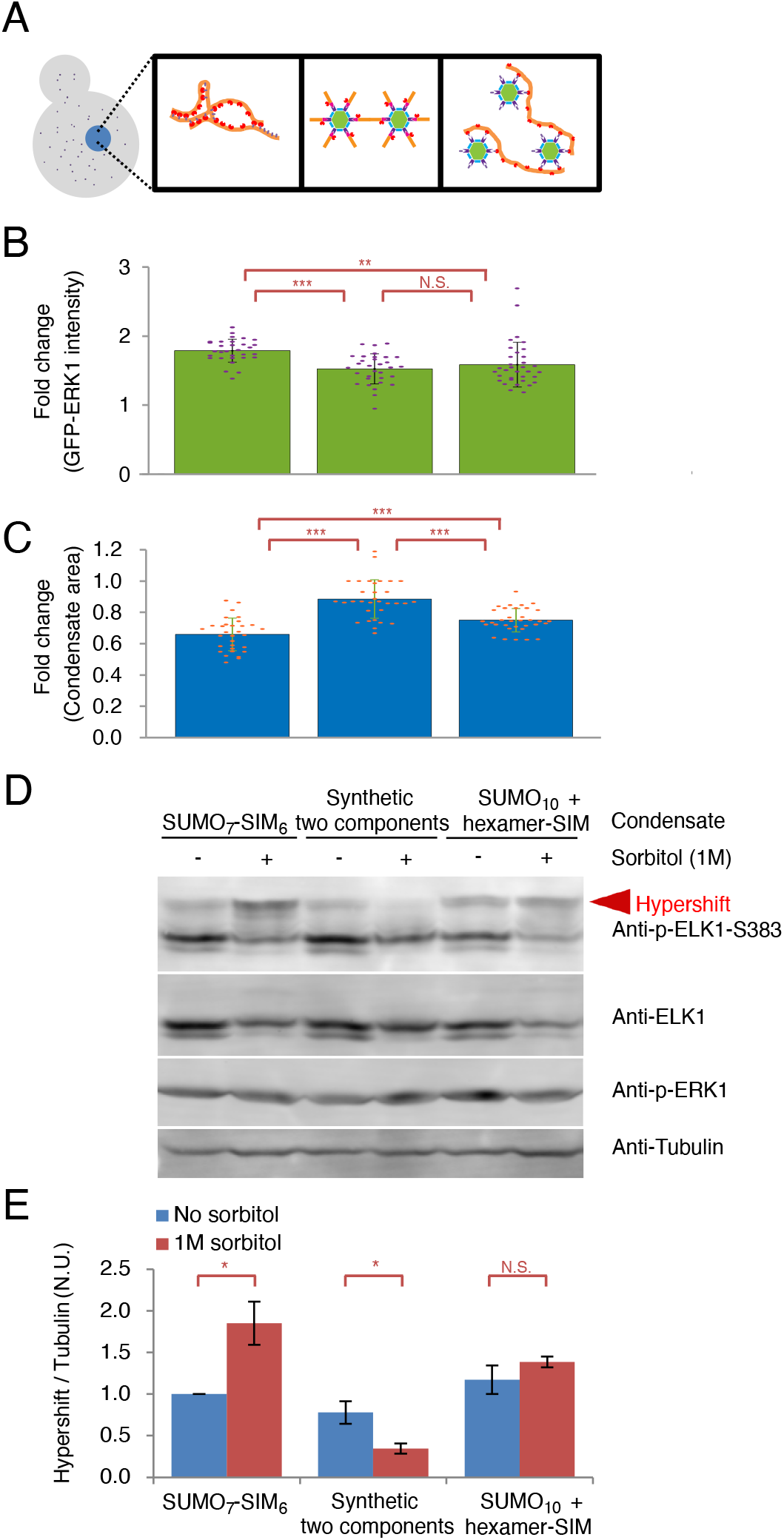
Synthetic condensed phase signaling can respond to osmotic compression. (A) Schematic showing three different types of synthetic condensate analyzed in this experiment: (Left) the relatively rigid two-component condensate; (middle) the flexible SUMO_7_-SIM_6_ condensate; (right) a mixed system with a SIM tagged hexameric component forming a network with a flexible SUMO_10_ chain. (B) GFP-ERK1-SIM concentration increases in condensates after osmotic compression. Average projections of z-stacks were used to measure the GFP intensity in condensates before and after 1h osmotic compression with 1M sorbitol. Bar graphs show mean, ± SD, n > 30. Statistical comparisons are by Student’s t-test: **p<0.01, 1***p<0.001. (C) Condensate volumes decrease to varying degrees upon osmotic compression, depending on scaffold flexibility. Cells were immobilized, and average projections of z-stacks were used to measure the area of condensates. The same condensate was measured before and after 1h osmotic compression with 1M sorbitol. Each point represents a condensate. Bar graphs show mean, ± SD, n > 30. Statistical comparisons are by Student’s t-test: ***p<0.001, N.S., not significant. (D) Representative western blots are shown for (from top to bottom) the ELK1-S383 phosphoepitope, total ELK1, the activation-loop phosphate on the ERK1 kinase (pT202/pY204), and tubulin loading control. Red arrowheads indicate hypershifted bands that are quantified below. (E) Osmotic compression increases hyperphosphorylation, only in condensates with flexible scaffold components. Quantification of degree of hyperphosphorylation from western blots. The intensity of hypershifted ELK1 bands were normalized to tubulin expression. The hyperphosphorylation level of ELK1 in the leftmost strain was set as 1. *hog1Δ* strains were used to prevent osmoadaptation. Error bars indicate ± SD, n = 3. Statistical comparisons are by Student’s t-test: *p<0.05, N.S., not significant.

When *HOG1* is deleted, crosstalk from osmotic stress pathway to the *Fus3* kinase has been reported (O’Rourke and Herskowitz, 1998). We demonstrated above that *Fus3* can phosphorylate ELK1 when recruited to condensates (Figure 2B). Therefore, it was important to rule out the possibility that somehow osmotic stress leads to activation of *Fus3* and its recruitment to condensates. To exclude this possibility, we also examined the ELK1 hyperphosphorylation response after osmotic compression in a *hog1Δ; fus3Δ* double mutant strain. As shown in Figure 4 Supplement 3A, ELK1 hyperphosphorylation increased similarly upon osmotic shock in *hog1Δ; fus3Δ* double mutant strains and *hog1Δ* strains. Therefore, *Fus3* does not appear to play a role in the change in hyperphosphorylation that we observe after osmotic compression.

In a control strain containing a soluble SUMO_2_ scaffold control without condensates, there was significant degradation of mCherry-ELK1-SIM (Figure 4 supplement 4A). There was less ELK1 degradation in all strains that recruited mCherry-ELK1-SIM to condensates, presumably because they are protected from proteolysis. ELK1 S383 phosphorylation was almost undetectable upon sorbitol treatment in SUMO_2_ scaffold control strain. Additionally, we tested a SUMO_2_ scaffold control in which SUMO was fused the coiled-coil domain from TRIM25. The levels of ELK1 S383 phosphorylation relative to total ELK1 protein were decreased by almost 80% in this soluble SUMO_2_ dimer scaffold strain (Figure 4 Supplement 4B). This decrease may be caused by reduced molecular diffusion upon the increase in macromolecular crowding.

The initial client concentration was higher in SUMO_10_-SIM_6_ condensates than in SUMO_7_-SIM_6_ condensates (Figure 4 supplement 1). This led to the hypothesis that the hyperphosphorylation is almost saturated in SUMO_10_-SIM_6_ condensates, while SUMO_7_-SIM_6_ condensates remain sensitive to concentration changes. To test this idea, we used stronger promoters to increase client concentration in SUMO_7_-SIM_6_ condensates. Consistent with the hypotheses, there was no significant change in ELK1 hyperphosphorylation upon osmotic compression in these strains (Figure 4 supplement 2). Therefore, condensed phase signaling can respond to osmotic compression when the system is appropriately tuned; a consistent activation of hyperphosphorylation is seen in SUMO_7_-SIM_6_ condensates that are basally relatively inefficient when they contain a low enough client concentration that reaction rates are not saturated.

Next we investigated the effect of osmotic compression on phosphorylation within the more rigid two-component condensates. We used the lower affinity Im2 variant that gives more liquid-like condensate, which we reasoned would be more likely to respond to compression (Heidenreich et al., 2020). Upon osmotic compression, condensate area was only slightly decreased (∼ 10%, Figure 4C), but the concentration of GFP-ERK1-SIM clients increased in the condensates to a similar degree as in SUMO_7_-SIM_6_ condensates, by ∼60% (Figure 4B). Strikingly, osmotic compression did not increase hyperphosphorylation in the two-component condensates, but rather the degree of ELK1 hyperphosphorylation was decreased by ∼50% upon osmotic compression (p < 0.05, Figure 4D and 4E). We screened for colonies with lower concentration of clients in two-component condensates to rule out the possibility that the phosphorylation rates were initially saturated. The degree of hyperphosphorylation in these strains was still reduced upon osmotic compression (Figure 4 supplement 5). Together, these results suggest that rigid condensates are relatively insensitive to osmotic compression, and suggest that the flexibility of SUMO-SIM condensates is important for their ability to transduce osmotic stress to changes in hyperphosphorylation.

To further investigate the hypothesis that structural rigidity explains the lack of response of the two-component condensates to osmotic compression, we modified this two-component system to connect each hexamer subunit with a flexible SUMO_10_ polypeptide. In place of the Im2-E9 interaction, we simply placed a SIM peptide at the C-terminus of the hexamer subunits, such that the network was crosslinked by SUMO-SIM interactions (Figure 4 Supplement 6A). Co-expression of the hexamer-SIM with SUMO_10_ led to condensate formation and client recruitment (Figure 4 Supplement 6B). The area of these condensates area was reduced by 25% upon sorbitol treatment, significantly more than that for rigid two component condensates (p < 0.001, Figure 4C and Figure 4 Supplement 7). This increase in condensate compression is consistent with the idea that polymer flexibility is important for mesoscale structural changes to synthetic condensates upon changes to the physical environment. Western blotting showed that the relative amount of ELK1 hyperphosphorylation in the hexamer-SIM + SUMO_10_ condensate slightly increased, but not significantly (Figure 4D and 4E). Therefore, this mixed system with one structured and one flexible component shows intermediate behavior. These results further support the hypothesis that condensate compressibility and scaffold linker flexibility are important for the response of condensed-phase phosphorylation to osmotic compression.

### Synthetic condensed phase signaling responds to changes in macromolecular crowding

We previously found that the mTORC1 pathway influenced the phase separation of SUMO_10_-SIM_6_ synthetic condensates through modulation of the concentration of ribosomes in the cytosol (Delarue et al., 2018). About 30-40% of the volume of the cytosol is taken up by macromolecules, and ribosomes constitute about half of this excluded volume (Delarue et al., 2018; Ellis and Minton, 2003a). We previously found that, when mTORC1 is inhibited by rapamycin, the concentration of ribosomes decreases 2-fold, and the phase separation of SUMO_10_-SIM_6_ is substantially reduced (Delarue et al., 2018) A similar effect is observed in the *sfp1Δ* mutant, which lacks the *Sfp1* transcription factor that drives high levels of ribosome biogenesis downstream of active mTORC1; this mutant has low baseline crowding and therefore low SUMO_10_-SIM_6_ phase separation, but crowding and condensation can be rescued by osmotic compression. We explored these conditions and mutants to test the hypothesis that condensed-phase signaling in SUMO_7_-SIM_6_ condensates respond to changes in macromolecular crowding (Figure 5A).

**Figure 5:**
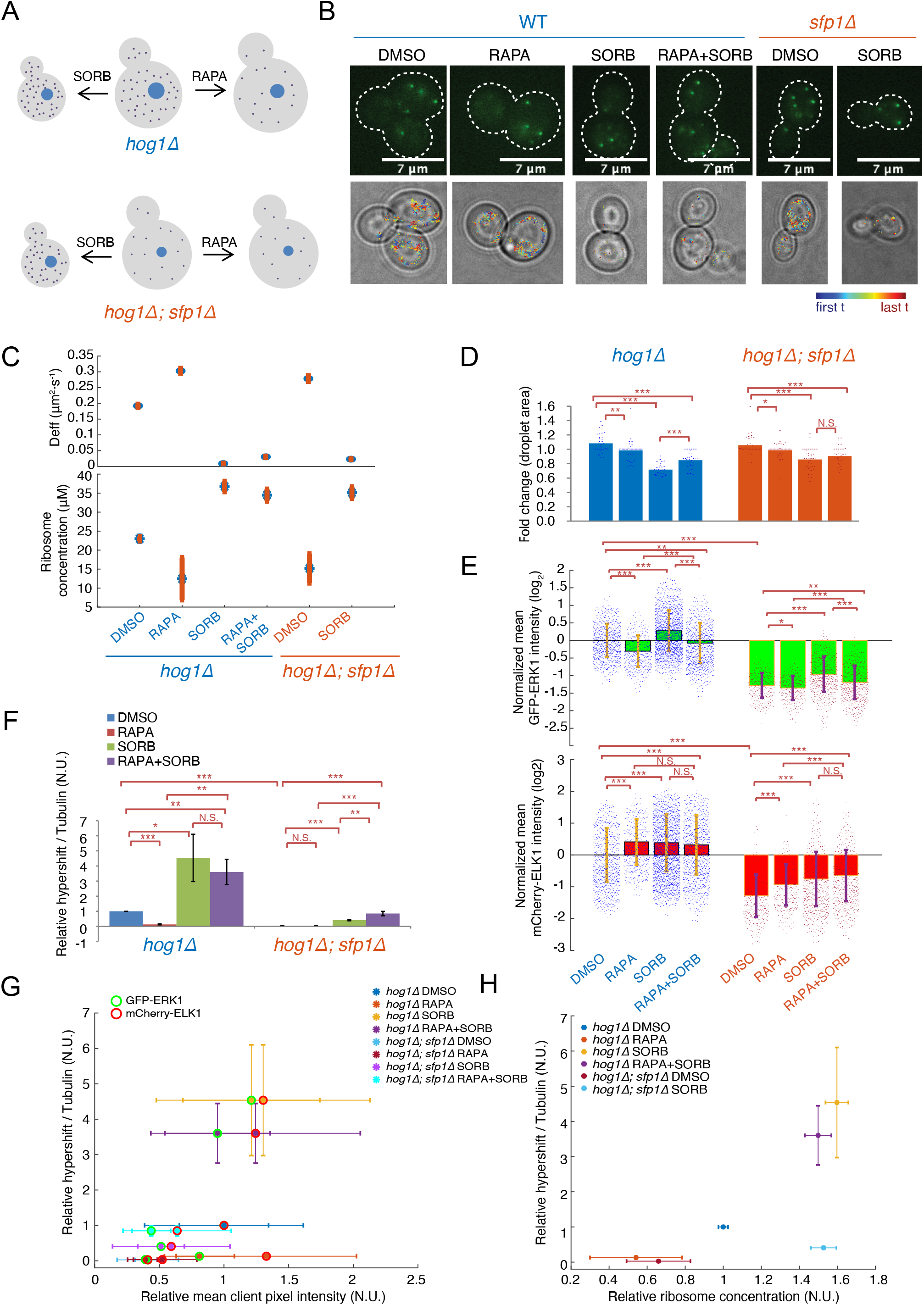
Synthetic condensed phase signaling responds to changes in macromolecular crowding. Schematic of experiments to assess the response of SUMO_7_-SIM_6_ condensates to changes in molecular crowding. *hog1Δ* (top) and *hog1Δ; sfp1Δ* mutant (bottom) yeast cells were treated with 1 μM rapamycin for 2 hours to inhibit mTORC1, which leads to decreased ribosome concentrations and decreased macromolecular crowding. The dots in the cells indicate molecular crowders such as ribosomes, that decrease in concentration in response to rapamycin in *hog1Δ* cells. Ribosome concentrations are constitutively low in *hog1Δ; sfp1Δ* cells and do not change in response to rapamycin. *hog1Δ* strains were used to prevent osmoadaptation. (B) Representative fluorescent images of cytosolic 40nm-GEM in *S. cerevisiae* under different conditions. GEM trajectories from movies (Supplementary Movie 2) were projected onto brightfield images, blue represents the start of trajectory while red indicates the trajectory end. Scale bar = 7*µm*. (C) Median effective diffusion coefficients D_eff_ of GEMs (top) as well as estimated ribosome concentrations (bottom) under different conditions, error bars represent standard error of mean. (D) SUMO_7_-SIM_6_ condensate area is reduced in response to rapamycin and osmotic compression in *hog1Δ* cells, but the rapamycin effect is attenuated in the *hog1Δ; sfp1Δ* mutant. Cells were treated with 1 μM rapamycin/DMSO for 2 hours, 1M sorbitol for 1 hour, or 1 μM rapamycin/DMSO for 1 hour followed by 1-hour treatment with both 1M sorbitol and 1 μM rapamyclin/DMSO. Cells were immobilized, and average projections of z-stacks were used to measure the area of condensates. The same condensate was measured before and after treatment, and its fold-change was plotted as a single point, n > 30. (E) Quantification of GFP-ERK1-SIM recruitment to SUMO_7_-SIM_6_ condensates in various crowding conditions. Concentrations of GFP-ERK1-SIM within SUMO_7_-SIM_6_ condensates were measured using fluorescence intensities from average projections of z-stacks. The concentrations were normalized to the median of the leftmost strain (*hog1Δ* with DMSO) and are plotted on a log_2_ scale. Each point represents a single condensate. Error bars indicate ± SD; statistics comparisons are by Student’s t-test: *p < 0.05; **p < 0.01; ***p < 0.001; N.S., not significant. (F) Reduction of ELK1 phosphorylation by rapamycin could be reversed by sorbitol treatment. To quantify ELK1 phosphorylation from western blots, band intensities of total Ser383 phosphorylated ELK1 were normalized to tubulin levels, and this value was further normalized to the phosphorylation level of ELK1 in the leftmost strain (*hog1Δ* with DMSO). All bar graphs show mean ± SD, n = 3; statistical comparisons are by Student’s t-test: *p < 0.05, ** p < 0.01, *** p < 0.001, N.S., not significant. (G) Scatter plot of relative hypershift / tubulin data from F were plotted against relative mean GFP-ERK1-SIM (green circle) and mCherry-ELK1-SIM (red circle) client pixel intensity from E. Different conditions were represented through different colors of lines and circle inside. Error bars are standard deviations. (H) Scatter plot of relative hypershift / tubulin data from F were plotted against relative ribosome concentration from C. Colors of lines and dots indicate conditions. Vertical error bars are standard deviations while horizontal error bars are standard error of means. All statistical comparisons are by Student’s t-test: *p<0.05, ***p<0.001, N.S., not significant.

**Figure 6:**
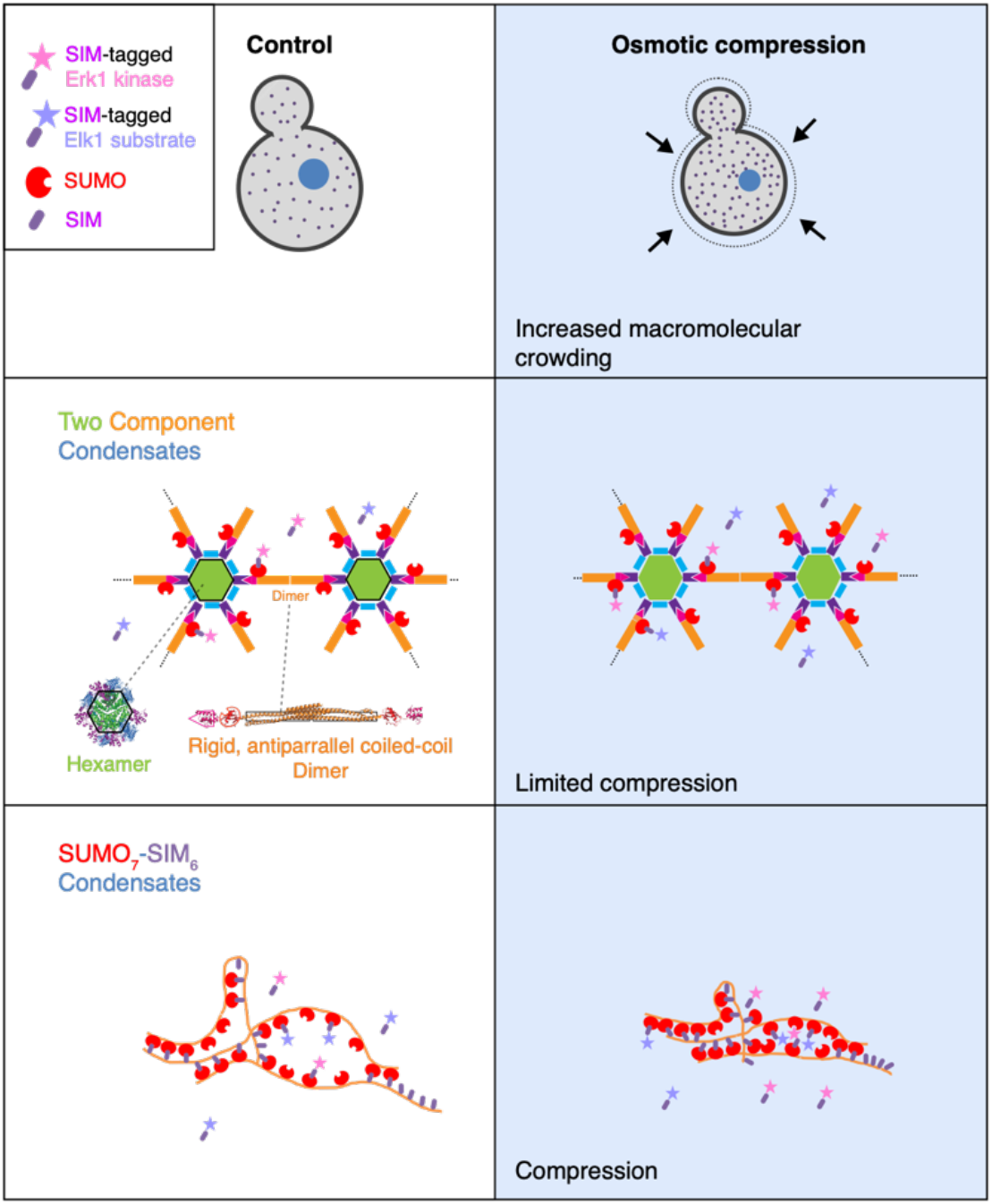
Proposed model for hyperphosphorylation in condensates under hyperosmotic compression. **(Top)** Schematic of the osmotic compression experiment. After addition of 1M sorbitol, cell volume decreases and macromolecular crowding increases. **(Middle)** In the synthetic two component condensate system, the bound clients are spaced out by a rigid scaffold that does not greatly potentiate phosphorylation. Increased macromolecular crowding upon osmotic compression does not greatly impact the organization of the rigid scaffold components of this condensate and therefore phosphorylation is not affected. **(Bottom)** In SUMO_7_-SIM_6_ condensates client proximity can vary substantially depending on the local conformation of the network of flexible SUMO_7_-SIM_6_ scaffold polymers. Upon osmotic stress, increased macromolecular crowding leads to compression of these condensates and therefore kinases and substrates are brought into closer proximity, leading to enhanced phosphorylation.

We took advantage of 40-nm diameter genetically encoded multimeric nanoparticles (40nm-GEMs) to quantify the degree of macromolecular crowding in cells (Delarue et al., 2018). 40nm-GEMs are comprised of a self-assembling scaffold (the encapsulin from *Pyrococcus furiosus*, PDB: 2E0Z (Akita et al., 2007)), fused to a fluorescent protein (mSapphire, (Zapata-Hommer and Griesbeck, 2003)). We imaged cells at 100 Hz to obtain tracks of the motion of 40nm-GEMs (Figure 5B, Supplemental Movie 2). From the effective diffusion coefficient 40nm-GEMs, we can infer properties of the intracellular environment, including the degree of macromolecular crowding at the length-scale of ribosomes. For example, we previously determined that the effective diffusion coefficient (D_eff_) was a reliable indicator of macromolecular crowding of the cytoplasm by ribosomes (Delarue et al., 2018). A fully experimentally parameterized version of the Doolittle equation can predict 40nm-GEM D_eff_ as a function of ribosome concentration and *vice versa* (Delarue et al., 2018; Doolittle, 1951). Consistent with previous results, we found that the baseline D_eff_ was 0.192 µm^2^ s^-1^ (predicting 23 µM ribosomes), and that this value increased to ∼0.303 µm^2^ s^-1^ upon 2 hours treatment with 1 µM rapamycin (Figure 5C). This increased D_eff_ is consistent with an almost 2-fold reduction of ribosome concentration to ∼13 µM (Figure 5D). The *sfp1Δ* mutant had a similar increase in 40nm-GEM D_eff_ to 0.3 µm^2^ s^-1^, indicating a similar almost 2-fold decrease in ribosome concentration, to 13 µM, but in this case without perturbation to mTORC1 kinase activity (Figure 5C, D). Osmotic compression of cells with 1M sorbitol for one hour led to an ∼10-fold decrease in 40nm-GEM D_eff_ in all conditions (Figure 5B and 5C), consistent with increased macromolecular crowding, to ∼ 35 µM ribosome concentration.

We previously showed that the size of SUMO_10_-SIM_6_ condensates was substantially reduced upon decrowding of the cytoplasm by mTORC1 inhibition or deletion of *SFP1*. We therefore expressed a higher concentration of SUMO_7_-SIM_6_ to favor condensate formation even when cytosolic crowding is reduced. We quantified the area of the same condensate in single *hog1Δ* cells before and after 2h treatment with DMSO (solvent control) or rapamycin, and found that inhibition of mTORC1 did not cause a significant decrease in condensate area (Figure 5D, Figure 5 supplement 1A), presumably because we are substantially above the phase boundary in these cells and so high macromolecular crowding is not required for condensation. Osmotic compression had a larger impact on condensate area. Similarly to Figure 4C, osmotic compression led to a roughly 30% area decrease, and this effect was partially offset when cells were pretreated with rapamycin; 1h rapamycin treatment followed by 1h sorbitol treatment resulted in a 20% average decrease in condensate area. In *sfp1Δ; hog1Δ* cells, which have reduced macromolecular crowding even without mTORC1 inhibition, condensate area was only slightly impacted by rapamycin treatment, while osmotic compression gave a 20% decrease in area. Macromolecular crowding could both impact the degree of phase separation of SUMO_7_-SIM_6_ and also compress the condensates. The most striking effect was that condensate area was substantially reduced when macromolecular crowding was increased. Increased macromolecular crowding is predicted to increase phase separation, and thus increase condensate area. The fact that condensate area is decreased upon osmotic compression suggests that increased macromolecular crowding actually compresses condensates.

The partitioning of clients into condensates could also respond to mcaromolecular crowding, either due to changes in the condensate *per se*, or due to changes in interactions between the clients and scaffolds (Nakashima et al., 2019) However, changes in macromolecular crowding and mTORC1 activity had modest effects on client concentration in condensates, and no consistent patterns were observed. In *hog1Δ* cells, mTORC1 inhibition with rapamycin only slightly decreased the average intensity of GFP-ERK1-SIM in condensates relative to DMSO control (∼25% decrease, p < 0.001, Figure 5E, top) upon rapamycin treatment, while mCherry-ELK1-SIM fluorescence slightly increased (∼ 25% increase, p < 0.001, Figure 5E, bottom). Western blotting showed that rapamycin treatment substantially reduced total ELK1 levels (∼70%, Figure 5F, Figure 5 supplement 1B), possibly due to the global translational inhibition effects of rapamycin treatment (Fingar et al., 2002). Osmotic compression with 1M sorbitol slightly increased the average intensity of GFP-ERK1-SIM (∼25%, p < 0.001), while again mCherry-ELK1-SIM fluorescence increase slightly (∼25%, p < 0.001). In *sfp1Δ; hog1Δ* cells, the average intensity of both mCherry-ELK1-SIM and GFP-ERK1-SIM were reduced ∼2-3 fold relative to *hog1Δ* cells, however Western blotting indicated that the total levels of ELK1 were substantially reduced, ∼4-to 5-fold (Figure 5 supplement 1B and 1C), again possibly due to the reduced translational capacity of these cells (Fingerman et al., 2003). Together, these results indicate that client partitioning into SUMO_7_-SIM_6_ condensates is not strongly impacted by changes in macromolecular crowding.

We next investigated how total phosphorylation and hyperphosphorylation in SUMO_7_-SIM_6_ condensates was affected by changes in macromolecular crowding. Figure 5F shows the hyperphosphorylation of ELK1, quantified from the fraction of slowly migrating band in the ELK1-phosphoS383 western blot relative to tubulin levels. This metric accounts for the changes in ELK1 concentration after rapamycin treatment and in *sfp1Δ; hog1Δ* cells. Hyperphosphorylation was reduced in conditions with decreased macromolecular crowding (e.g. 10-fold decease in *sfp1Δ; hog1Δ* cells relative to *hog1Δ* cells) and increased in conditions with increased macromolecular crowding (e.g. >10-fold increase after osmotic compression in both *hog1Δ* and *sfp1Δ; hog1Δ* cells.)

Changes to macromolecular crowding have complex effects on ELK1 total concentration, client partitioning, condensate area, and hyperphosphorylation. In principle, changes in the concentration of kinase and substrate within condensates could partly explain the changes in hyperphosphorylation by simple mass action. However, we found no clear relationship between client concentration in condensates and fraction hyperphosphorylation (Figure 5G), arguing against this simple model. However, when we plotted fraction hyperphosphorylation against ribosome concentration, a clear trend was apparent. (Figure 5H). High ribosome concentration correlated with a high degree of hyperphosphorylation of ELK1, and low ribosome concentration correlated with low hyperphosphorylation. The *hog1Δ; sfp1Δ* strain under osmotic compression was an apparent outlier to the trend, but both ERK1 and ELK1 concentrations are around 3-fold lower in condensates in this strain, likely accounting for the lower than expected hyperphosphorylation when normalized to tubulin (there is still a 10-fold increase in hyperphosphorylation at high ribosome concentration, but the baseline in *hog1Δ; sfp1Δ* strains is very low). If we calculated hyperphosphorylation normalized to total phosphoS383-ELK1, or if we simply quantify total S383 phosphorylation, this outlier is resolved and the trend holds for all conditions (Figure 5, supplement 3). No matter which metric we choose, there is no clear correlation with client concentration, but the correlation between ribosome concentration and Elk1 phosphorylation is very clear. Therefore, we conclude that condensed phase signaling within SUMO_7_-SIM_6_ condensates responds to macromolecular crowding.

## Discussion

### Condensates facilitate phosphoregulatory network rewiring

It has been suggested that biomolecular condensates could act as signaling hubs to integrate biological information. PML-nuclear bodies, the biological inspiration for the synthetic polySUMO-polySIM condensates used in this study, are an example. Here, SIM motifs in the HIPK2 kinase and its substrate P53, lead to recruitment to PML-nuclear bodies. P53 is then phosphorylated, leading to induction of apoptosis (Sung et al., 2011). PML-nuclear bodies also recruit multiple other kinases including ATR (Barr et al., 2003). Indeed, many kinases have now been reported to localize to condensates (Zhang et al., 2021a), or be recruited into condensates under stress (Reineke and Lloyd, 2015; Shattuck et al., 2019; Takahara and Maeda, 2012; Wippich et al., 2013; Zhang et al., 2021a). However, it has been very difficult to dissect the physiological relevance of this organization because mutations that disrupt condensation could have pleiotropic effects. Our synthetic approach was designed to circumvent this issue.

Using our synthetic system, we made a number of observations that suggest that new kinase-substrate connections can be generated more easily within condensates than in solution. We were able to drive strong phosphorylation of the ELK1 substrate with non-cognate kinases Fus3 and Cdk1, simply through co-recruitment to a synthetic condensate (Figure 2). In both cases, dynamic phosphorylation changes were observed as kinase activity was modulated. Furthermore, we found that recruitment to condensates allowed non-cognate substrates and non-consensus phosphoacceptor motifs to be phosphorylated by multiple different kinases (Figure 2; Supplements 5 and 6). This expansion of specificity could have interesting ramifications for biology. It will be interesting to investigate whether the activity and substrate specificity of kinases that are recruited to endogenous condensates is altered. The consensus sequences that are often used to predict kinase substrates may be less important in the context of condensed phase signaling and a larger number of possible phosphorylation sites may need to be investigated. For example, it has been shown that the crucial regulatory sites of some kinase substrates are actually at non-canonical sites; for example, degradation of the yeast cell cycle regulator Sic1 is triggered by multisite phosphorylation including non-consensus Cdk1 sites (Koivomagi et al., 2011; Nash et al., 2001). By extension, kinase-substrate interactions within condensates could lead to unexpected modes of phosphoregulation.

A further prediction of the ease with which we generated dynamic phosphorylation within condensates is that recruitment to condensates may facilitate the evolution of new links in phosphoregulatory networks. Phosphorylation can occur in condensates even in the absence of any obvious docking or consensus sites. It is possible that these initial phosphorylation events could provide a starting point from which useful regulation could evolve.

### Beyond mass-action, condensate flexibility and high-densities of client binding sites are important for efficient condensed-phase signaling

Recently a number of studies have reported the accelerated biochemical activities in condensates (Huang et al., 2019; Peeples and Rosen, 2021; Poudyal et al., 2019). A recent study (Peeples and Rosen, 2021) found that reactions were accelerated by mass action. In addition, they found that certain scaffolds decreased the effective K_M_ of the reaction, suggesting that molecular organization was important for strong activity enhancement. In this study, we used two distinct synthetic condensates, one based on a flexible scaffold polymer (polySUMO-polySIM) and another based on a highly structured network (the hexamer, dimer system) and varied the number of free client binding sites, the affinity and valency of client-scaffold interactions, client expression levels, the material properties of condensates, and condensate flexibility (Figure 3). Many of these factors modulated client concentration within condensates, and this somewhat correlated with the degree of hyperphosphorylation of ELK1. However, we took advantage of multiple designs of synthetic condensate to further investigate the hypothesis that specific organization of clients by the condensate scaffold are important for activity enhancement in our systems. Consistent with the idea that client proximity within condensates is important, we found that a well-ordered, relatively inflexible hexamer-dimer condensate with widely spaced client binging sites (SUMO domains) was quite inefficient compared to the flexible polySUMO-polySIM condensates. Furthermore, the main factor that had a strong effect on hyperphosphorylation beyond the correlation with client concentration was the number of excess client binding sites; SUMO_10_-SIM_6_ condensates gave much better activity enhancement than SUMO_7_-SIM_6_ condensates, even when the latter condensates had higher client concentration. Strikingly, the presence of up to triple-SUMO and high client concentration within rigid networks was insufficient to drive strong hyperphosphorylation. Therefore, we speculate that dense clusters of >3 SUMO domains within the flexible SUMO-SIM condensates can. organize clients within. dense pockets that enable multisite phosphorylation.

The design of our hexamer-dimer condensates allowed us to vary scaffold interaction strength by modulating Im2-E9 binding affinity (K_d_) from ∼10^−5^ M to ∼10^−8^ M. This large change in scaffold dynamics had a relatively modest effect, increasing hyperphosphorylation ∼2-fold (Figure 3E). Modulating scaffold binding affinity in these condensates was previously shown to change material properties from liquid-like to solid-like (Heidenreich et al., 2020). However, the diffusion of scaffold proteins and clients are quite different inside condensates (Banani et al., 2016). Diffusion of SIM-tagged client in our experiments, unlike scaffold molecules, may be relatively unaffected by Im2-E9 affinity change, as their diffusion is mainly determined by their interaction with SUMO, which is independent of the Im2-E9 interaction. Our attempts to characterize client-diffusion by Fluorescence Recovery After Photobleaching (FRAP) were unsuccessful, because client diffusion was very rapid and the condensates are small (typically ∼ 1 µm diameter). With stronger interactions between client and scaffold, the diffusion of scaffolds within the condensed-phase is likely to become more important. However, in our system, we speculate that condensed-phase phosphorylation is robust to large changes in scaffold dynamics (i.e. four-orders of magnitude change in interaction strength) because clients can diffuse rapidly through a fluid phase between scaffold molecules within the condensates.

In conclusion, we found that client concentration is correlated with the efficiency of condensed-phase signaling, but mass action is insufficient to explain activity enhancement; condensate flexibility and the ability to form local high-density clusters of client binding sites are crucial factors.

### Synthetic condensed-phase signaling can respond to biophysical changes

Recently, we found that macromolecular crowding strongly affects biomolecular condensation(Delarue et al., 2018). As stated above there are a large number of signaling molecules, including kinases that localize to condensates (Zhang et al., 2021a). Furthermore, several recent examples of endogenous condensates have now been reported to respond to macromolecular crowding (Cai et al., 2019). This leads to the hypothesis that there is an axis of control spanning from the global biophysical state of the cell, to the degree of phase separation and the material properties of condensates, and finally to molecular-scale biochemical reactions. However, it has been extremely difficult to investigate the idea using traditional reductionist approaches or manipulation of endogenous molecules. Here, we were able to directly investigate this hypothesis using kinases in synthetic condensates.

Osmotic compression leads to water efflux in cell, resulting in a reduced cell volume, increased macromolecular crowding and decreased molecular diffusion (Joyner et al., 2016; Miermont et al., 2013). For the condensates that we used in this study, we found that condensate size was reduced by osmotic compression to a degree that depended on the scaffold protein flexibility (Figure 4C). Strikingly, we found that the degree of hyperphosphorylation in SUMO_7_-SIM_6_ condensates was increased ∼2-fold by osmotic compression. By perturbing ribosome concentration, we determined that this osmotic response was due to changes in macromolecular crowding (Figure 5). In contrast, the more rigid hexamer-dimer condensates did not significantly compress and failed to transduce changes in macromolecular crowding, and neither the SUMO_10_-SIM_6_ nor SUMO_7_-SIM_6_ condensates with high client concentration responded to osmotic compression, presumably because hyperphosphorylation was already nearly saturated (Figure 4). Therefore, phosphorylation reactions within condensates can either respond to changes in biophysical parameters such as macromolecular crowding, or be insensitive to these changes depending on their properties and the parameters of the system. The ability of condensed-phase chemical reactions to respond to macromolecular crowding presents exciting new possibilities for both synthetic biology and the elucidation of new mechanisms of biological regulation and homeostasis. For example, the mechanisms that sense mechanical compression remain poorly understood (Delarue et al., 2018), but mechanical compression leads to increases in macromolecular crowding and we have now demonstrated that the macromolecular crowding can modulate phosphorylation rates within condensates. It will be exciting to investigate whether kinases in endogenous condensates can transduce mechanical information through these biophysical mechanisms.

## Materials and methods

### Yeast transformation

Yeast strains were created by transforming with a LiAc based approach according to Cold Spring Harbor Protocols. All strains were constructed in the W303 strain background (*MATa leu2-3, 112 trip1-1 can1-100 ura3-1 ade2-1 ade2-1 his3-11-,15*). A list of strains built is provided in Table S1. To tag Cdk1 and Fus3 with GFP-SIM, pFA6a-CDK-GFP-SIM or pFA6a-FUS3-GFP-2xSIM plasmid was cut within the CDS of Cdk1 or Fus3, and the linearized plasmid was then transformed into W303.

### Plasmid construction

The open reading frames encoding SUMO_10_-SIM_6_ condensates, GFP-ERK1-1SIM and mCherry-ELK1-1SIM were chemically synthesized (Qinglan, China). The SUMO_10_-SIM_6_ condensate expression plasmid in yeast was constructed by fusion 5’ end of the ORF with the strong promoter from *TDH3* by Gibson assembly into the pRS306 vector (Sikorski and Hieter, 1989). The kinase expression plasmid in yeast was constructed by fusion of the 5’ end of the ORF with yeast weak promoter (Pab1) or medium promoter (Rpl18) by Gibson assembly (Gibson et al., 2009) into the pAV105 vector (Agmon et al., 2015). The substrate expression plasmid in yeast was similarly constructed by fusion with yeast weak promoter (Rnr2) or medium promoter (rpl18) and assembled into the pAV103 vector (Agmon et al., 2015). To express the synthetic condensate component in yeast, the dimer and hexamer component were amplified by PCR and assembled into pRS306 vector or pRS304 vector (Sikorski and Hieter, 1989) by Gibson assembly. The WW reporter was generated by fusion of GFP, 2xWW (WW domain from Peptidyl-prolyl cis-trans isomerase NIMA-interacting 1, *Homo sapiens*) with Rpl18 promoter and assembled into the pRS304 vector. To tag Cdk1 and Fus3 with GFP-SIM, Cdk1/Fus3 CDS, GFP and SIM tag were amplified and assembled into pFa6a vector by Gibson (Bahler et al., 1998). All yeast plasmids were integrated into the host genome. pET28b vectors were used for bacterial expression. Open Reading Frames (ORFs) of the SUMO_10_, FKBP12-SIM_6_, FRB-mCherry-ERK1 and FRB-GFP-ELK1 proteins were fused N terminally to the 6X histidine tag for purification. The ORFs and the vectors were Gibson assembled. A list of plasmids constructed is provided in Table S2.

### Protein purification

Protein purification from *E. coli* cells was performed as previously described (<u>Howard et al., 2014</u>). Briefly, proteins were expressed in Rosetta2 DE3 competent cells by induction with 100 μM IPTG for 18 hr at 16°C. Bacterial culture were collected and centrifuged at 4000 rpm for 20 min at 4°C. The cell pellet was resuspended in cold lysis buffer (50 mM NaH2PO4, 300 mM NaCl, 10 mM imidazole pH7.6, 1 mM PMSF). After sonication, the lysate was centrifuged at 12000 rpm for 20 min at 4°C. The supernatant was mixed with magnetic Ni-NTA beads (Qiagen) and incubated for 2 hr at 4°C. The bound beads were collected and rinsed 3 × with wash buffer containing (50 mM NaH2PO4, 300 mM NaCl, 20 mM imidazole pH7.6). The bound proteins were eluted with elution buffer (50 mM NaH2PO4, 300 mM NaCl, 500 mM imidazole pH7.6). The eluted proteins were concentrated using Amicon® Ultra Centrifugal Filters (Millipore Sigma). The concentrated proteins were dialyzed into SUMO-SIM protein buffer (150 mM KCl, 20 mM HEPES pH 7, 1 mM MgCl2, 1 mM EGTA, 1 mM DTT) with 10% glycerol using PD10 columns (GE Healthcare), followed by further concentration using an Amicon Ultra 30K device (Millipore) at 4C., and finally flash frozen in aliquots with liquid nitrogen and stored at -80°C.

### In vitro kinase assays

For phosphorylation in condensates, the components for kinase reaction and condensate formation were divided into two halves, each with a 10 µl volume. Mixture of two the halves initiated kinase reaction. The first half contained 30 µM FKBP12-SIM_6_, 40 nM FRB-GFP-ERK1, 8 µM FRB-mCherry-ELK1, 50 µM Rapamycin (or same volume of DMSO), 10 mM MgCl2 and 75 mM KCl. Components were equilibrated for 20 minutes. The phosphorylation was started by addition of the second half, which contained 24 µM SUMO_10_, 200 µM ATP, 10 mM MgCl2, 75 mM KCl and 0.1 µCi of [γ-32P]ATP. Two halves were mixed thoroughly by gentle pipetting. Reactions were carried out at room temperature for 5 minutes and terminated by addition of 10 µl 5x SDS loading buffer (10% SDS, 0.5 M DTT, 50% glycerol, 0.25% Bromophenol blue). All samples were separated on 4–12% Bis-Tris gels (ThermoFisher Scientific). The gel was dried and exposed to a phosphor screen. Phosphor screens were analyzed with a Typhoon 9500 scanner (GE) using ImageQuant software (GE).

### ERK1-GFP partition coefficient calculation

While the reaction proceeded, the other half of the solution was transferred to a 384-well plass bottomiamging plate and imaged using an Andor Yokogawa CSU-X confocal spinning disc on a Nikon TI Eclipse microscope and fluorescence was recorded with a sCMOS Prime95B camera (Photometrics) with a 100x objective (pixel size: 0.11um). A stack of 50 images were acquired at 1µM intervals from the bottom of the plate upwards. In order to determine the camera background value, a blank well with buffer was imaged. This background value was subtracted from every image in order to allow for more accurate partition coefficient analysis The fluorescence integrated density of ‘drops’ or ‘solvent’ in the 488 and 561 channels was calculated using a classifier mask derived from contrast adjusted imaged segmented with the trainable Weka Segmentation package on default settings. The condensates are heavier than the solution and sediment near the bottom of the well. Therefore, the reaction is asymmetrically distributed along the z axis and symmetric along the x & y axes. We reasoned that in order to calculate the true partition coefficient of a given volume, it would be necessary to calculate the total protein in the solvent and condensates along the entire vertical stack. It was calculated that given the volume of reaction added to the well, there would be an additional 2490 ‘slices’ spaced out every 1uM on top of the 50uM already imaged. It was observed that condensates never formed in slices 48-51 as the condensates had sedimented beyond this point. It should also be noted that carefully controlling temperature and evaporation are required to prevent solution turbidity. Therefore, the average solvent protein concentration was determined by finding the mean integrated density per slice in this ‘top of the well’ equivalent from slices 48-51. Next, the condensate and solvent protein concentrations across the entire well were calculated by integrating along the slices using the ‘trapz’ function from the pcrma package (Borchers and Borchers, 2021). In order to calculate partition coefficient of the reaction, the total protein in condensates was therefore divided by the total concentration in solution across the entire y dimension. The scripts used for partition coefficient calculation are provided in supplementary files as QC1 and QC2.

### Western blots

Cell cultures were grown to OD 0.6–0.8. Cells were collected and treated with 1M LiAc for 5 min on ice. After centrifugation cell pellets were treated with 0.4M NaOH for 5 min. Then cells were centrifuged again, and were suspended in SDS-PAGE sample buffer and boiled for 5min. Supernatant samples were separated on 10% SDS-PAGE gels and transferred to Immobilon-FL PVDF membrane (Millipore). Membranes were probed with primary antibodies, including anti-p-ELK1-S383, anti-ELK1, anti-p-ERK1, anti-Tubulin, anti-HA, anti-p-P53-S37, anti-p-P53-S9, anti-p-RPS6-S235/236 and anti-p-Sata3-Y705 (Key resources table in STAR methods), and corresponding secondary IRDye 800CW antibodies (LI-COR Biosciences). Band intensities were quantified with Image Studio™ analysis software (LI-COR Biosciences).

### Cell treatments for western blot

For western blot with alpha factor treatment, cells at log phase (OD around 0.8) were treated with 2 µM for 1h. Then cells were collected by centrifugation for 3min at 3000g and lysed for western blot.

For western blot with sorbitol treatment only, cells at log phase (OD around 0.8) were collected by centrifugation for 3min at 3000g. Cells were next suspended in synthetic complete with dextrose medium (SCM) with or without 1 M sorbitol and grown for 1h. Then cells were collected by centrifugation for 3min at 3000g for western blot.

For western blots with rapamycin treatment, cells at log phase (OD around 0.8) were incubated with DMSO (solvent control) or 1 µM rapamycin for 2h. Then cells were collected as above for western blot. For treatment with rapamycin combined with sorbitol, cells were first incubated with 1 µM rapamycin for 1 h, then collected by centrifugation for 3min at 3000g. Cells were next suspended in SCM with 1 µM rapamycin and 1 M sorbitol for 1 h. Then cells were collected for lysis and SDS-PAGE followed by western blotting. For the control experiments to rapamycin or rapamycin plus sorbitol, same volume of DMSO was used instead.

### Phosphatase treatment of yeast lysates

Yeast cell were collected when the culture OD reach 0.8. Cells were pretreated with 0.4M NaOH for 2 min on ice. After centrifugation, cell pellets were suspended in lysis buffer with protease inhibitor tablet (Pierce) and lysed by bead beating. Then samples were centrifuged at 800g for 2 minutes. The supernatants were used with lambda protein phosphatase (NEB) treatment according to its protocol. For treatment with phosphatase inhibitor, NaF (final 50 mM), β-glycerol phosphate (final 50 mM), and Na3VO4 (final 1 mM) were added. The lysates were incubated at 30°C for 30 minutes and terminated by addition of 5x SDS loading buffer followed by boiling for 5 minutes.

### Imaging and quantification of fluorescence intensity inside condensate and condensate size

Cells were imaged using TIRF Nikon TI Eclipse microscope in epifluorescence mode, and fluorescence was recorded with an sCMOS camera (Zyla, Andor) with a 100x objective. GFP and mCherry channel images were generated by average projection of 13 z-slices with 0.4 μm spacing (4.8 μm total). Condensate properties within cells were characterized using ImageJ plugin, TrackMate (Tinevez et al., 2017). Due to relatively higher intensity and higher contrast in GFP fluorescent signals, condensates in GFP channel were first detected using LoG (Laplacian of Gaussian filter) detector with one micron ‘Estimated blob diameter’ and fixed ‘Threshold’ cross different experimental conditions. For each individual image with both GFP and mCherry channels, number of detected condensates within GFP channel was then recorded and was used as a criterion for choosing the ‘Threshold’ parameter for condensate detection in mCherry channel. ‘Estimated blob diameter’ parameter for mCherry condensate detection still maintained as one. Using this method, the majority of condensates detected in both channels overlapped with each other. By saving the particle detection results from Trackmate as *xml* files, we then extracted and compiled particle properties in both channels, especially condensate mean pixel intensity, using home-written Matlab code. In addition, we also extracted the background mean pixel intensity by randomly selecting 20 circles in each image from areas away from cellular condensates. Thus, the final condensate mean pixel intensity were calculated by subtracting the background mean pixel intensity from condensate mean pixel intensity identified above.

For calculation of condensate area and client concentration changes, log-phase cells were immobilized in 384-well imaging plates coated with concanavalin A (ConA). GFP and mCherry channel images were generated by average projection of 13 z-slices with 0.4 μm spacing (4.8 μm total). For experiments with sorbitol treatment only, immobilized cells were imaged, then media was carefully removed and replaced with media containing 1M sorbitol, and the same cells were imaged again after 1h. For experiments under various conditions, such as rapamycin treatment, cells were immobilized, imaged and then media was carefully changed, and cells were imaged again after 2h with 1 µM rapamycin. For treatment with rapamycin and sorbitol, cells were imaged, then were treated sequentially with 1µM rapamycin for 1h, then old medium was removed and incubated with fresh SCM with 1µM rapamycin and 1M sorbitol for a further 1h, and cells were imaged again. Condensates were identified manually based on their GFP signal, and the area of the same condensate before and after treatment was calculated by measurement in Image J.

### HILO imaging of GEMs

GEM particles in *S. cerevisiae* yeast cells were imaged using Highly inclined thin illumination (HILO) TIRF Nikon TI Eclipse microscope in partial TIRF mode under 100% power of 488nm excitation laser. The emitted fluorescent signals were transmitted through a 100x objective (100x Phase, Nikon, oil NA = 1.4, part number = MRD31901) and recorded with a sCMOS camera (Zyla, Andor, part number = ZYLA-4.2p-CL10). The GFP filter set (ET-EGFP (FITC/Cy2), Chroma, part number = 49002) was embedded within the light path, which includes an excitation filter (Excitation wavelength/ Bandwidth (FWHM) = 470/40 nm), a dichroic mirror (long pass beamsplitter, reflecting < 495 nm and transmitting > 495 nm wavelength) and an emission filter (Emission wavelength/ Bandwidth (FWHM) = 525/50 nm). Each GEM movie was recorded at a single focal plane with 10ms frame rate for total 4 s, using Nikon NIS-Elements Advanced Research software.

### Calculation of effective diffusion constant

For every 2D GEM trajectory, we calculated the time-averaged mean-square displacement (MSD) at different time intervals:

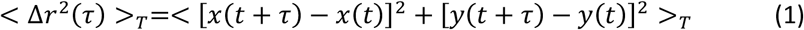

where ‘<>_*T*_’ represents time averaging for each trajectory of all displacements under time interval τ

To reduce tracking error due to particles moving in and out of focal plane, we limited our analyses to particle trajectories with longer than 10 time points. Time-averaged MSD for each trajectory is then fitted using a linear time dependence at first 10 time intervals:

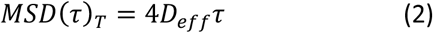

where *D*_*eff*_ is effective diffusion coefficient for each trajectory, with the unit of *µm*^2^/*s*.

For each experimental condition, we then used median value of *D*_*eff*_ among all trajectories and plotted bar graph with error bars as standard error of mean for characterizing GEM mobility.

### Relative ribosome concentration calculation

Based on the phenomenological Doolittle equation (Doolittle, 1951) as well as a derived model from Delarue et al (Delarue et al., 2018), which describes the relationship between cellular crowding due to macromolecules such as ribosomes and the effective diffusion coefficients of mesoscale tracer particles.

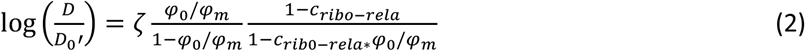

where D represents the experimental effective diffusion coefficient, D_0_’ represents the effective diffusion coefficient under control condition, *ϕ*_0_ is the volume fraction of macromolecules under control condition, *ϕ*_*m*_is the maximum volume fraction of macromolecules when there is no water, *ζ*is the dimensionless interaction parameter, *C*_*ribo − rela*_ is the relative ribosomal concentration compared to control condition.

## Acknowledgements

We thank Michael Rosen, Allyson Rice and members of the HHMI Summer Institute at the Marine Biology Institute in Woods Hole for discussions that initiated this project. We thank Jef Boeke, David Engelberg, Meta Heidenreich and Emmanuel D. Levy for sharing yeast strains and plasmids, Greg Brittingham for help with imaging and analysis. We thank Lance Denes and Srinjoy Sil for critical reading of the manuscript. We thank the rest of the Holt lab for helpful discussions. This work was funded by NIH R01 GM132447 and R37 CA240765, the American Cancer Society, the Pershing Square Sohn Cancer Award, the Chan Zuckerberg Initiative, and the Air Force Office of Scientific Research (AFoSR grant FA9550-21-1-3503 0091).

## Notes

### Competing Interest Statement

The authors have declared no competing interest.

