## Supplemental Figures for "Synthetic Condensed-Phase Signaling Expands Kinase Specificity and Responds to Macromolecular Crowding"

##### Figure 1; Figure Supplement 1

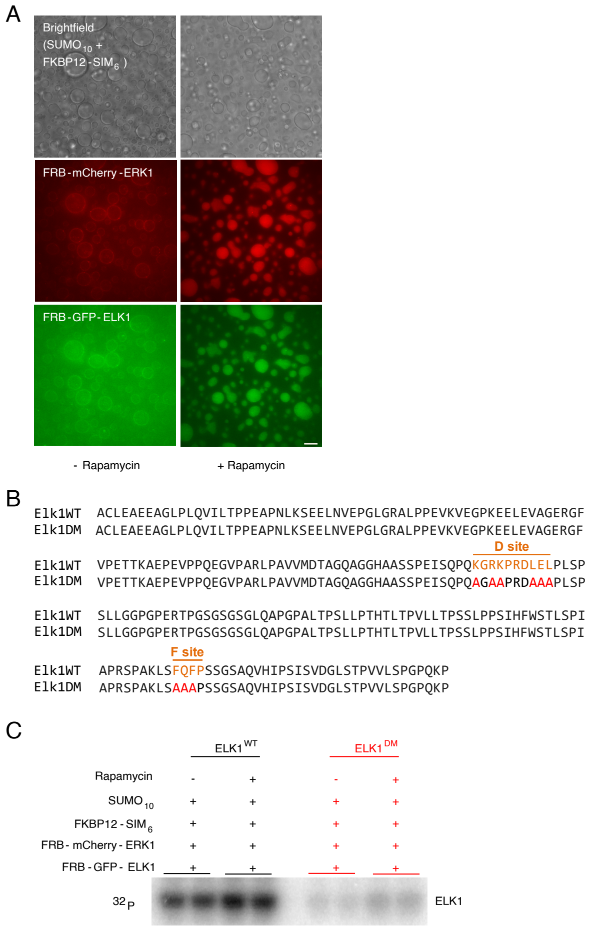


**Figure 1 Supplement 1: Recruitment of kinase and substrate into synthetic condensates increased phosphorylation *in vitro***

(A) Micrographs showing rapamycin-induced client recruitment into condensates *in vitro*. Top, brightfield shows liquid condensates formed from $SUMO_{10}$ + FKMP12-$SIM_6$; middle shows FRB-ELK1-mCherry; bottom shows FRB-ERK1-GFP. Condensates were formed by mixing purified recombinant $SUMO_{10}$ and FKBP12-$SIM_6$ protein. Left column is control, right column is after client recruitment with rapamycin. Scale bar = 10 µm. (B) Amino acid sequences of $ELK1^{WT}$ and ELK1 docking site mutant peptides ($ELK1^{DM}$). The D and F docking motifs in $ELK1^{WT}$ are colored in orange. The mutated residues in $ELK1^{DM}$ are shown in red. (C) *In vitro* kinase assay data for Figure 1B. Kinase reactions were performed with and without rapamycin. $ELK1^{WT}$ and $ELK1^{DM}$ were used as substrates.

#### Figure 1; Figure Supplement 2

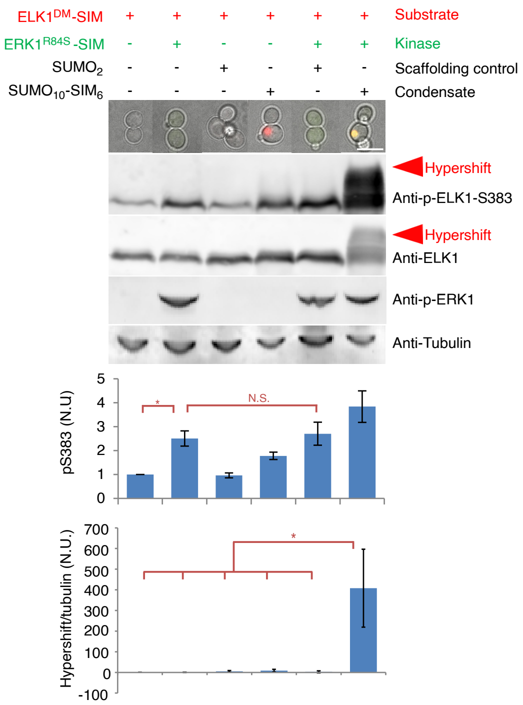


**Figure 1; Supplement 2: Characterization of ELK1$^{WT}$ phosphorylation in condensates *in vivo*.** Top panel, micrographs of control *S. cerevisiae* cells with various combinations of ERK1$^{WT}$ (green), ELK1 (red) and scaffold controls or $SUMO_{10}$-$SIM_6$ condensates (no fluorescent tag). SIM tags enabled client recruitment into $SUMO_{10}$-$SIM_6$ condensates. Scale bar = 5 µm. Below, representative Western blots for (from top to bottom) the ELK1-S383 phosphoepitope, total ELK1, the activation-loop phosphate on the ERK1 kinase (pT202/pY204), and tubulin loading control. Red arrowheads indicate hypershifted bands. For S383 phosphorylation, band intensities of total phosphorylated ELK1 were normalized to total ELK1 levels, and this value was further normalized to the phosphorylation level of ELK1 in the control strain (leftmost) expressing only ELK1. For hyperphosphorylation (Hypershift / Tubulin) quantification, the intensity signal of hypershifted band (detected by anti-p-ELK1-S383) was normalized to tubulin levels in the bottom graph, and the values were further normalized to the leftmost strain expressing ELK1 only. Error bars indicate ± SD, n = 3, Statistical comparisons are by Student's t-test: *p<0.05, N.S., not significant.

**Figure 1; Figure Supplement 3**

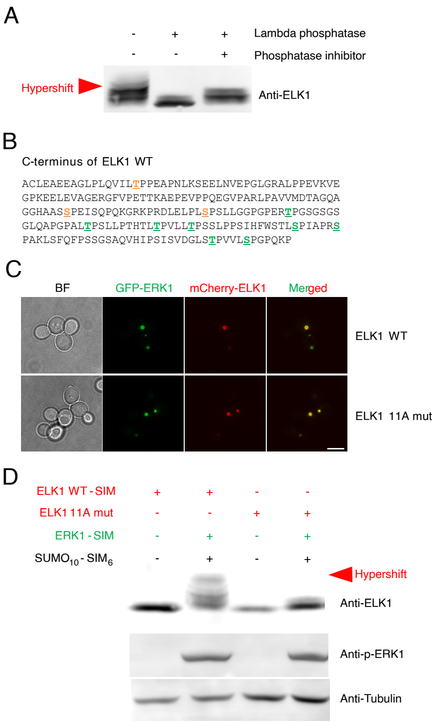


**Figure 1; Supplement 3: Recruitment to synthetic condensates leads to multi-site phosphorylation of ELK1 *in vivo***

(A) Western blot for total ELK1 of cell lysates treated with lambda phosphatase, in the absence or presence of phosphatase inhibitor. Red arrowhead indicate hypershifted bands. (B) Sequence of the C-terminus of ELK1. Known ERK1 phosphorylation sites are underlined in green, additional ERK1 consensus motifs (S/T-P) are underlined in orange. (C) Images showing client recruitment into condensates in cell. Clients were tagged with SIM. ERK1 was fused with GFP and the substrate, ELK1 or ELK1-11A mutant was fused with mCherry. Scale bar = 5 µm. (D) Representative western blots for total ELK1 (from top to bottom), the activation-loop phosphates on the ERK1 kinase (pT202/pY204), and tubulin loading control. Red arrowhead indicates the hypershifted bands that are not present in the ELK1-11A mutant.

#### Figure 2; Figure Supplement 1

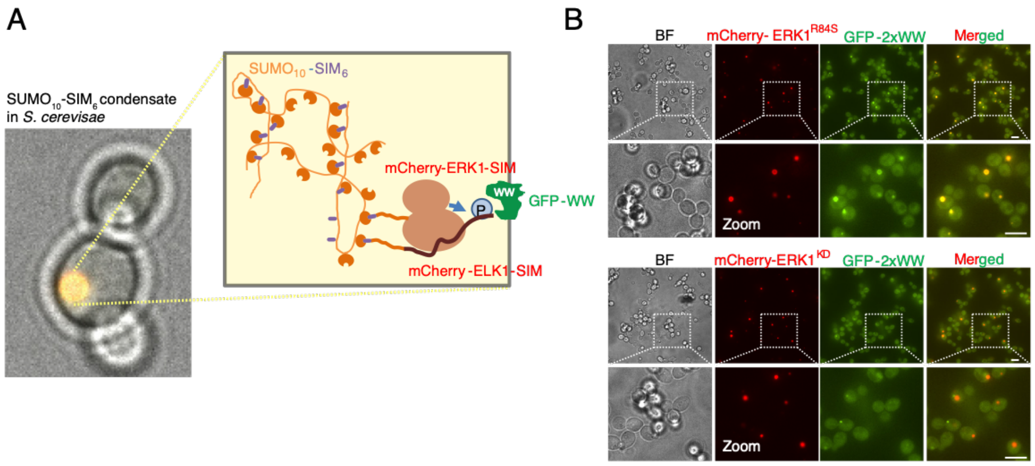


**Figure 2; Supplement 1: Validation of phosphorylation reporter using ERK1 as kinase**

(A) Schematic of the phosphorylation reporter comprised of a tandem repeat of the phosphopeptide binding WW domain from human PIN1 fused to GFP (GFP-2xWW). Upon kinase activation, the reporter binds to phosphorylated ELK1 in SUMO$_{10}$-SIM$_6$ condensates. ERK1 and ELK1 were tagged with SIM for recruitment into condensates and to mCherry for visualization. (B) Top two panels, co-expression of constitutively active mCherry-ERK1-3xSIM and mCherry-ELK1-SIM led recruitment of the GFP-WW reporter into condensates. Bottom, when a kinase dead (ERK1-K71R) mCherry-ERK1-3xSIM mutant (ERK1$^{KD}$) was used, the GFP-WW reporter did not colocalize to condensates, but rather localized to a separate punctate cellular structure (possibly spindle pole bodies), scale bar = 5 µm.

#### Figure 2; Figure Supplement 2

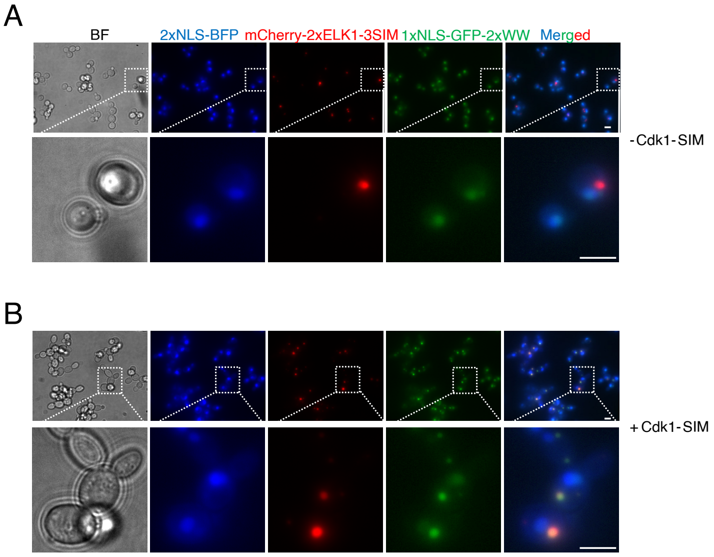


**Figure 2; Supplement 2: Validation of phosphorylation reporter using Cdk1 as kinase.** The basic design of phosphorylation reporter was slightly modified from the above system in Fig. 2, supp 1 to add competition between nuclear localization and condensate localization. The GFP-2xWW reporter was N-terminally tagged with an SV40 nuclear localization signal (NLS-GFP-2xWW). The endogenous copy of the cyclin dependent kinase Cdk1 was tagged at its C-terminus with a SIM tag to enable recruitment into $SUMO_{10}$-$SIM_6$ condensates. The mCherry-ELK1-SIM substrate was expanded to a tandem repeat to provide more potential phosphorylation binding sites for the reporter, and a 3xSIM tag was attached to its C-terminus (mCherry-2xELK1-3xSIM). The position of the nucleus was determined using a blue fluorescent protein (BFP) tagged at the N-terminus with two SV40 nuclear localization signals (2xNLS-BFP). (A) In a strain with no SIM tag on Cdk1 (top panels), the reporter is nuclear and there is no detectable GFP-WW signal in condensates. In contrast (B), in strains where Cdk1-SIM was targeted to condensates, in most cells, most of the GFP-WW reporter is in the condensates, and little signal is detected in the nucleus. GFP and mCherry channel images represent the average-intensity z-projection of a 4.8 µm z-stack with 0.4 µm step. Scale bars = 5 µm.

#### Figure 2; Figure Supplement 3

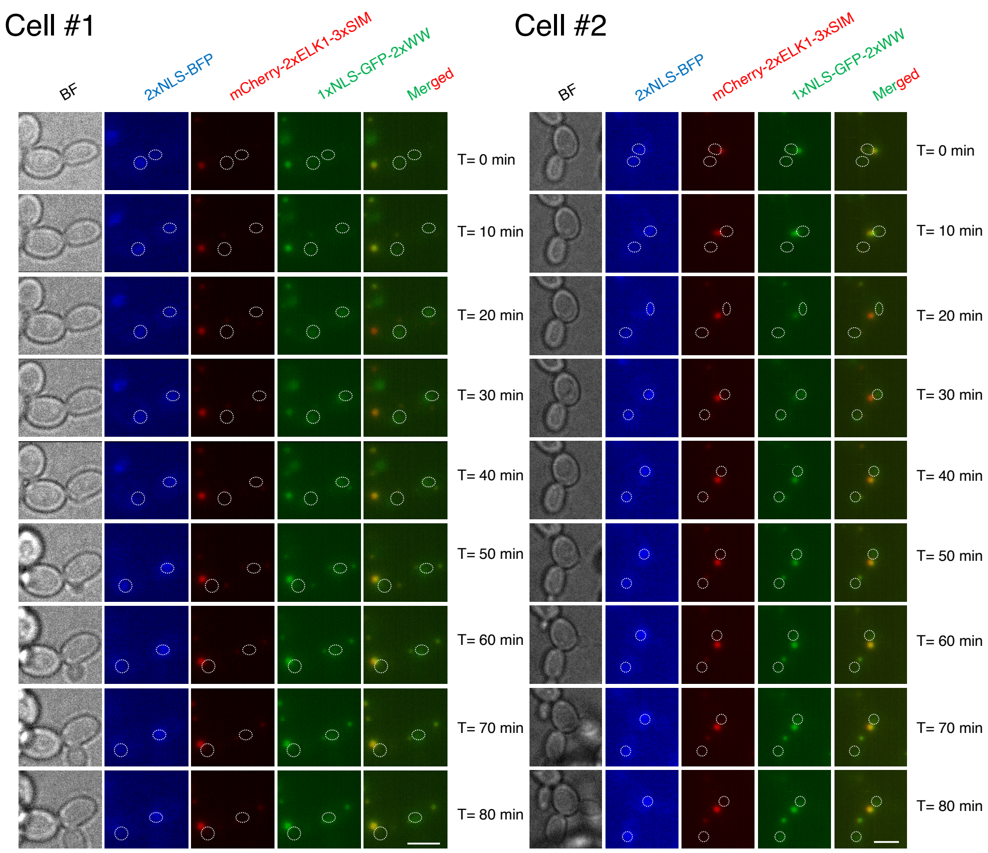


**Figure 2; Supplement 3: The NLS-GFP-2xWW phosphorylation reporter dynamically changes localization during mitotic exit.** Each row of micrographs shows an individual timepoint from movies of two mitotic *S. cerevisiae* cells. The point at which nuclear division initiated was set as t = 0. The channels are: brightfield (BF, grey); a reporter of nucleus position (2xNLS-BFP, blue); the ELK1 substrate (mCherry-2xELK1-3xSIM, red); and the phosphorylation reporter (NLS-GFP-2xWW, green). In the merged image, a white outline indicates the position of the nucleus. GFP and mCherry channel images are average-intensity z-projection of a 4.8 µm z-stack with 0.4 µm step. Scale bar = 5 µm.

#### Figure 2; Figure Supplement 4

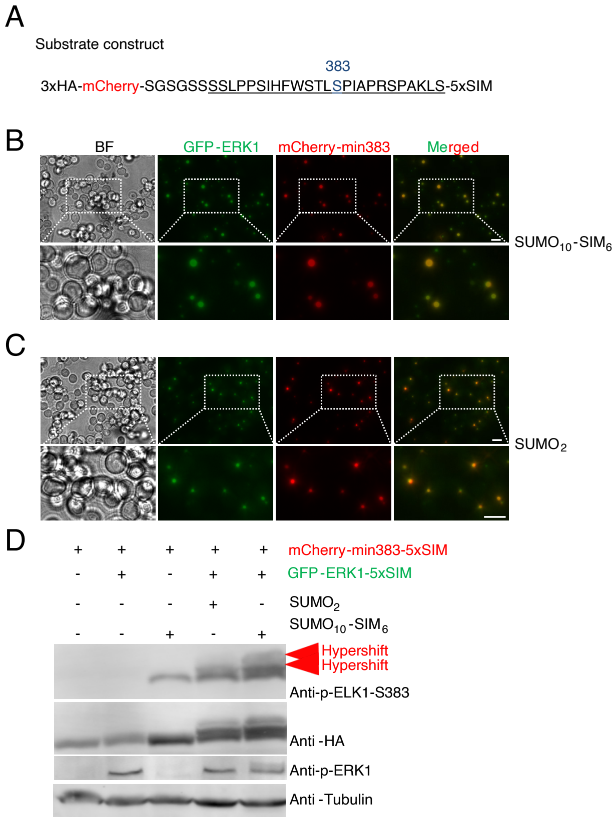


**Figure 2; Supplement 4: A minimal ELK1 peptide (min383) with docking sites completely removed is phosphorylated in condensates.**

(A) A minimal construct was designed that contains the S383 phosphoepitope from the C terminus of ELK1 but completely lacks all D site and F site docking motifs. The ELK1 sequence is underlined with S383 highlighted in blue. This peptide was fused to a triple HA epitope (3xHA) and mCherry at the N-terminus, and was tagged with five SIM tags at the C-terminus (5xSIM). (B) The min383 substrate is recruited to $SUMO_{10}$-$SIM_6$ condensates, but also forms condensates with $SUMO_2$ dimers. A 5xSIM tag gave robust recruitment to $SUMO_{10}$-$SIM_6$ condensates. Co-expression with $SUMO_2$ also resulted in small condensates (C), probably due to the multivalency of the 5xSIM tag. Scale bar = 5 µm. (D) The min383 substrate is more efficiently phosphorylated in both 2xSUMO and $SUMO_{10}$-$SIM_6$ condensates. Western results showing (from top to bottom) the ELK1-S383 phosphoepitope; anti-HA to indicate the total amount of min383 substrate; the activation-loop phosphate on the ERK1 kinase (pT202/pY204); and tubulin loading control. Red arrowheads indicate hypershifted bands.

#### Figure 2; Figure Supplement 5

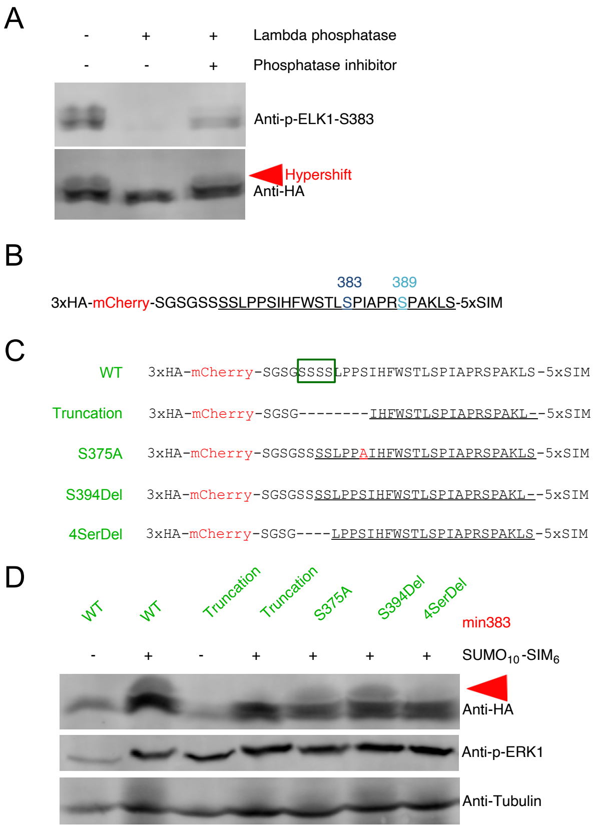


**Figure 2; Supplement 5: The hypershifted bands for the min383 substrate are largely due to four non-consensus serines**

(A) The hypershifted bands for the min383 substrate are due to phosphorylation. Cell lysates were treated with lambda phosphatase in the presence or absence of phosphatase inhibitor. Western results showing the ELK1-S383 phosphoepitope (top) and anti-HA to indicate the total amount of min383 substrate (bottom). Red arrowhead indicates hypershift bands. (B) Sequence of the min383 substrate. Two MAP kinase consensus motif serines (383 and 389) (serine followed by proline) are indicated in blue. (C) Constructs used for mapping the residues causing hypershift bands upon phosphorylation. Several point mutations or truncations were generated to identify the residues responsible for hypershifted bands. The four consecutive serines near the N-terminus are indicated in green box. (D) Western results showing (from top to bottom) the ELK1-S383 phosphoepitope; anti-HA to indicate the total amount of min383 substrate; the activation-loop phosphate on the ERK1 kinase (pT202/pY204); and tubulin loading control. Red arrowheads indicate hypershifted bands.

### Figure 2; Figure Supplement 6

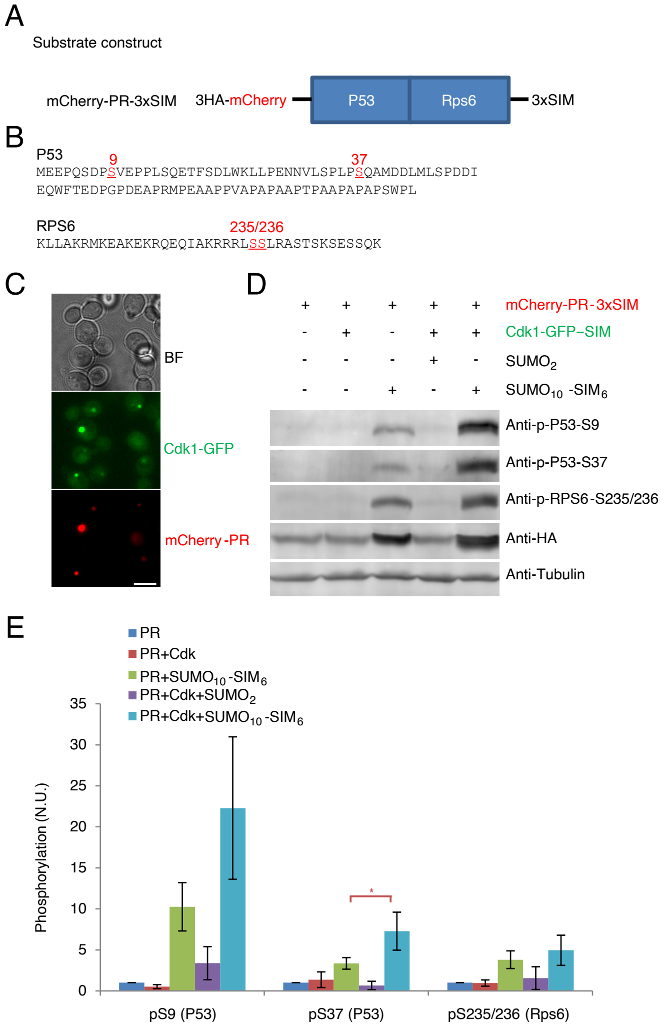


**Figure 2; Supplement 6: Cdk1 can phosphorylate non-consensus serines in a synthetic substrate (PR) when targeted to condensates.**

(A) Schematic of a synthetic substrate fusing short fragments from P53 and RPS6. A 3xHA tag and mCherry were fused to the N-terminus and a 3xSIM tag was attached to the C-terminus for recruitment into $SUMO_{10}$-$SIM_6$ condensates (mCherry-PR). (B) Sequences of the fragments from P53 and RPS6 used in the synthetic substrate. These sequences were chosen because reliable antibodies were available to detect the phosphoepitopes. The non-consensus serines tested for phosphorylation are underlined in red. (C) The mCherry-PR substrate and Cdk1 kinase clients were recruited into $SUMO_{10}$-$SIM_6$ condensates. Micrographs show that the kinase Cdk1 (green) and substrate PR (red), fused with SIM and 3xSIM respectively, colocalize to $SUMO_{10}$-$SIM_6$ condensates, bar = 5 µm. (D) The levels of phosphorylation of non-Cdk1-consensus sites are increased by co-recruitment into $SUMO_{10}$-$SIM_6$ condensates. Western blot results showing (from top to bottom) the P53-S9 phosphoepitope; the P53-S37 phosphoepitope; the RPS6-S235/S236 phosphoepitope; anti-HA to indicate the total amount of the mCherry-PR substrate; and tubulin loading control. (E) Quantification of Western Blots: For each quantification, total phosphorylated PR band intensities were normalized to total PR levels (anti-HA), and this value was then compared to the phosphorylation level of the strain expressing PR alone (leftmost), which was normalized to 1. Error bars indicate ± SD, n = 3. Statistical comparisons are by Student's t-test: *$p<0.05$.

#### Figure 3; Figure Supplement 1

| | | | | | | | | |
|---|---|---|---|---|---|---|---|---|
| ++ | + | ++ | + | ++ | ++ | ++ | ++ | SIM $K_d$ |
| 1 | 1 | 1 | 1 | 2 | 1 | 1 | 1 | Client #SIM |
| 4 | 4 | 4 | 4 | 4 | 4 | 1 | 1 | Excess SUMO |
| ++ | ++ | + | +++ | ++ | +++ | ++ | +++ | ELK1 expression |
| ++ | ++ | ++ | ++ | ++ | +++ | ++ | +++ | ERK1 expression |

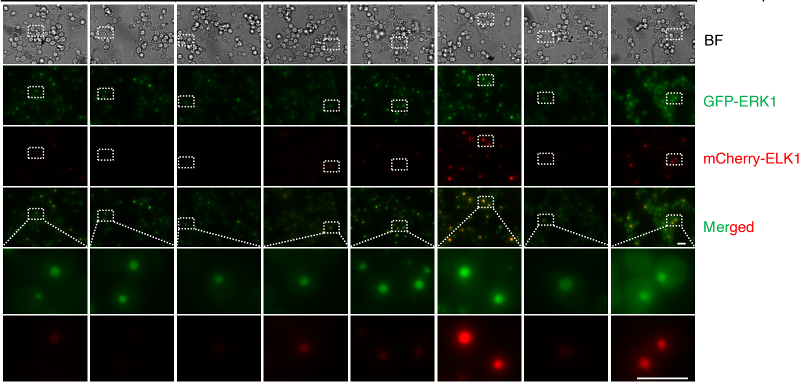


**Figure 3; Supplement 1: Representative images of client recruitment into SUMO-SIM condensates with varying client and scaffold properties.** Images showing recruitment of GFP-ERK1-SIM (or 2xSIM) and mCherry-ELK1-SIM (or 2xSIM) client recruitment into $SUMO_{10}$-$SIM_6$ or $SUMO_7$-$SIM_6$ condensates in *S. cerevisiae* cells. Mutant SIM motifs were also explored, and client expression levels were varied. Conditions for each experiment are summarized in the table above the micrographs. GFP and mCherry channel images represent average-intensity z-projections of 4.8 µm z-stacks with 0.4 µm step-size. Scale bars = 5 µm.

**Figure 3; Figure Supplement 2**

| | | | | | | | | | |
|---|---|---|---|---|---|---|---|---|---|
| ++ | ++ | + | ++ | + | ++ | ++ | ++ | ++ | SIM $K_d$ |
| 1 | 1 | 1 | 1 | 1 | 2 | 1 | 1 | 1 | Client #SIM |
| No condensate | 4 | 4 | 4 | 4 | 4 | 4 | 1 | 1 | Excess SUMO |
| ++ | ++ | ++ | + | +++ | ++ | +++ | ++ | +++ | ELK1 expression |
| ++ | ++ | ++ | ++ | ++ | ++ | +++ | ++ | +++ | ERK1 expression |

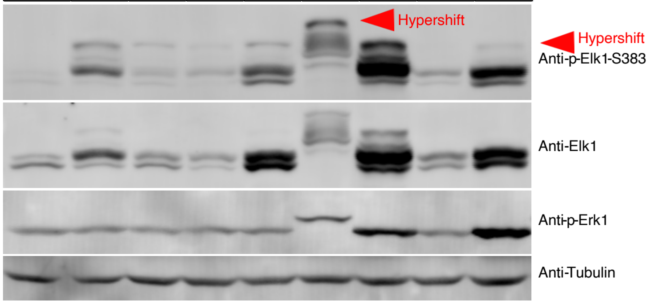


**Figure 3; Supplement 2: Representative western blot results showing ELK1 hyperphosphorylation as client and condensate properties are varied.** GFP-ERK1-SIM (or 2xSIM) and mCherry-ELK1-SIM (or 2xSIM) clients were recruited into $SUMO_{10}$-$SIM_6$ or $SUMO_7$-$SIM_6$ condensates in *S. cerevisiae* cells. Mutant SIM motifs were also explored, and client expression levels were varied. Conditions for each experiment are summarized in the table above the western blots. Representative western blots are shown for (from top to bottom) the ELK1-S383 phosphoepitope, total ELK1, the activation-loop phosphate on the ERK1 kinase (pT202/pY204), and tubulin loading control. Red arrowheads indicate hypershifted bands that were quantified in Figure 3. The 2xSIM tag leads to a slightly reduced gel mobility in the 6$^{th}$ lane from the left.

#### Figure 3; Figure Supplement 3

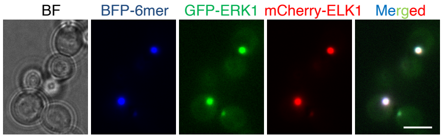


**Figure 3; Supplement 3: GFP-ERK1-SIM and mCherry-ELK1-SIM clients are efficiently recruited to the two-component condensates**. The hexamer-forming component was fused with blue fluorescent protein (BFP). Micrographs show representative average-intensity z-projections of BFP, GFP and mCherry channels. Z-stacks were 4.8 with 0.4 µm step-size. Scale bar = 5 µm.

#### Figure 3; Figure Supplement 4

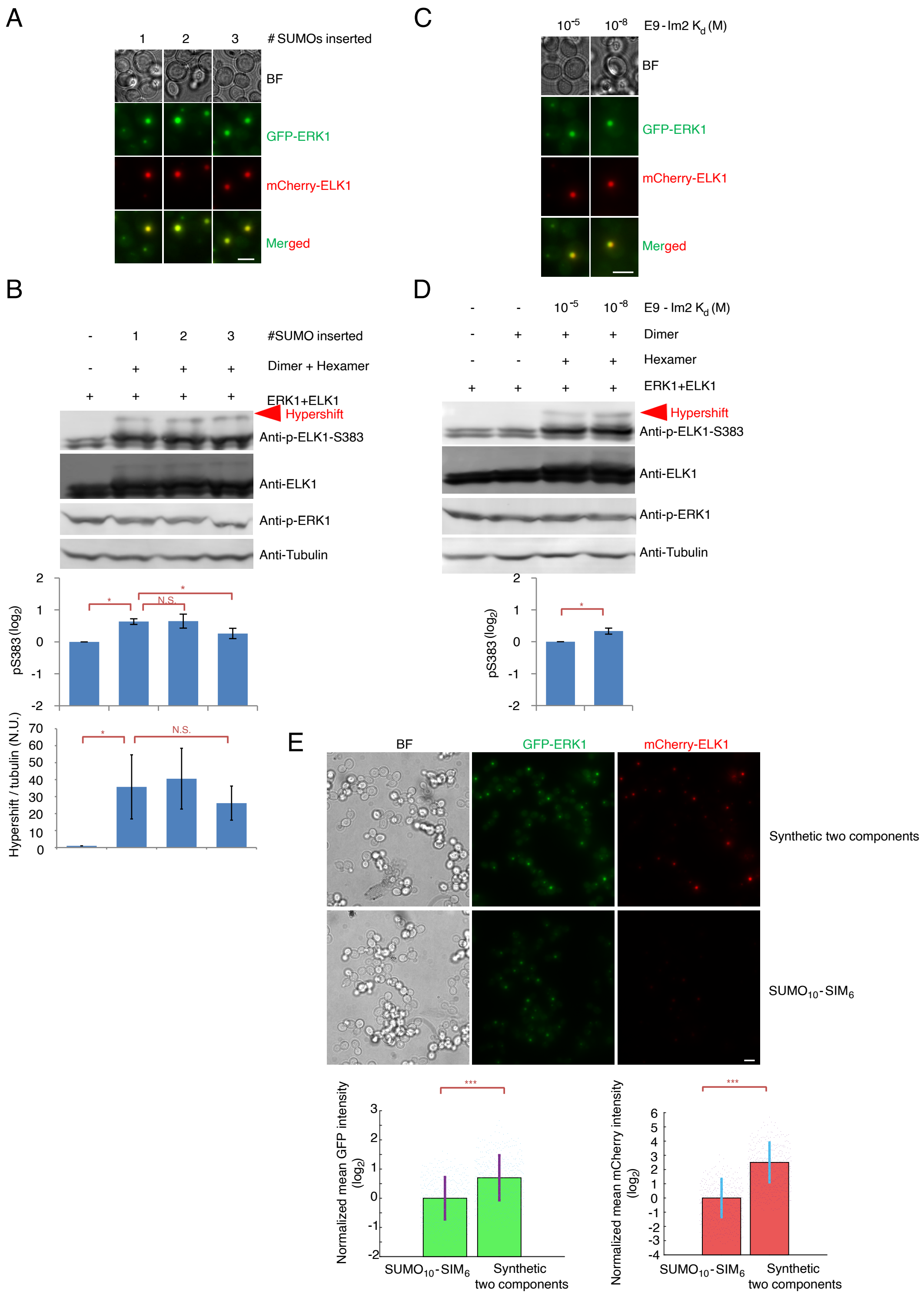


**Figure 3; Supplement 4: A relatively rigid two-component condensate also recruits clients and enhances phosphorylation but to a lesser degree than the flexible SUMO-SIM condensates.** (A) The two-component condensates recruit similar amounts GFP-ERK1-SIM and mCherry-ELK1-SIM clients with one, two or three SUMO domains per dimer-forming component. The hexamer-forming component was fused with blue fluorescent protein (BFP). Micrographs showing representative average-intensity z-projections of GFP and mCherry channels. Z-stacks were 4.8 with 0.4 µm step-size. Scale bar = 5 µm. (B) The two-component condensates drive similar amounts of ELK1 hyperphosphorylation with one, two or three SUMO domains per dimer-forming component. Representative western blots are shown for (from top to bottom) the ELK1-S383 phosphoepitope, total ELK1, the activation-loop phosphate on the ERK1 kinase (pT202/pY204), and tubulin loading control. Red arrowheads indicate hypershifted bands that were quantified in figure 3. For S383 phosphorylation, band intensities of total phosphorylated ELK1 were normalized to total ELK1 levels, and this value was further normalized to the phosphorylation level of ELK1 in the control strain (left) expressing only ELK1 and ERK1. The percentage of hypershifted bands relative to total S383 signal in ELK1 was quantified in the bottom graph. Error bars indicate ± SD, n = 3. Statistical comparisons are by Student's t-test: *$p<0.05$, N.S., not significant. (C) The two-component condensates recruit similar amounts GFP-ERK1-SIM and mCherry-ELK1-SIM clients with low and high Im2/E9 affinity. The dimer-forming component was fused with two Im2 variants, one with a micromolar affinity, the other with nanomolar affinity for E9. (D) Representative western blots are shown for (from top to bottom) the ELK1-S383 phosphoepitope, total ELK1, the activation-loop phosphate on the ERK1 kinase (pT202/pY204), and tubulin loading control. Red arrowheads indicate hypershifted bands that were quantified in figure 3. For Ser383 phosphorylation, band intensities of total phosphorylated ELK1 were normalized to total ELK1 levels, and this value was further normalized to the phosphorylation level of ELK1 in the control strain (left) with micromolar affinity to E9. Error bars indicate ± SD, n = 3. Statistical comparisons are by Student's t-test: *$p<0.05$. (E) The concentration of clients is higher in two component condensates than in $SUMO_{10}$-$SIM_6$ condensates. Top shows representative micrographs of *S. cerevisiae* cells expressing GFP-ERK1-SIM and mCherry-ELK1-SIM, and co-expressing either synthetic two-component condensates (single SUMO domain per dimer-forming component, and nanomolar Im2/E9 affinity, top panels) or a $SUMO_{10}$-$SIM_6$ condensate. The relative concentrations of ELK1 and ELK1 within condensates were estimated based on fluorescence intensities, normalized to the median value, and are shown in the green and red bar graphs, where each point is the quantification of a condensate. Error bars indicate ± SD. Statistical comparisons are by Student's t-test: ***$p<0.001$. .

#### Figure 4; Figure Supplement 1

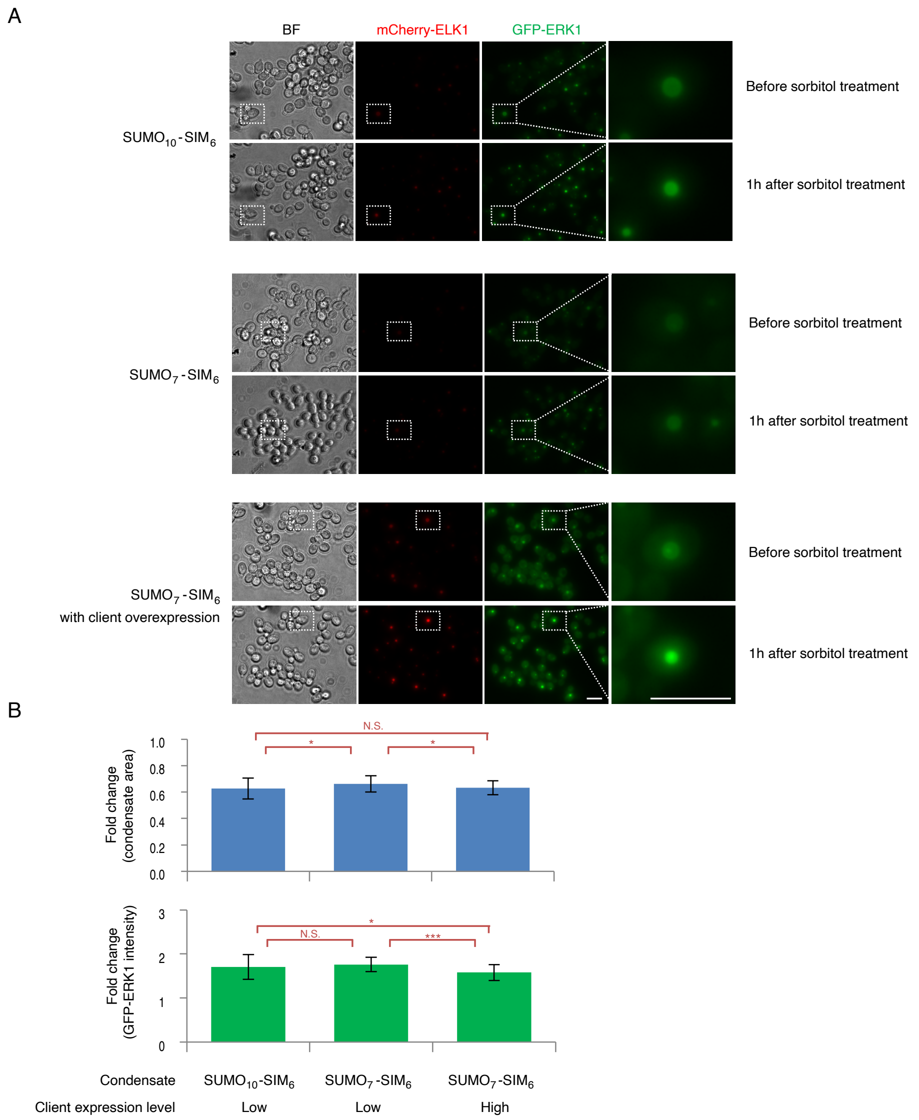


**Figure 4; Supplement 1: Osmotic compression leads to reduced condensate size and increased client concentration.**

(A) The reduction of condensate area and client concentration increase after 1 hour of osmotic compression with 1M sorbitol treatment were similar for $SUMO_{10}$-$SIM_6$ (top), $SUMO_7$-$SIM_6$ condensates (middle) and $SUMO_7$-$SIM_6$ with higher client expression levels (bottom). Cells were immobilized and the same cells were imaged before and after 1 hour of osmotic compression with 1M sorbitol. Representative micrographs are shown. Condensates were measured in average projections of z stacks that were 4.8 µm with 0.4 µm spacing. Scale bar = 5 µm. (B) Quantification of the change in condensate area and mean client concentration in the same condensates after 1h osmotic compression with 1M sorbitol. *hog1Δ* strains were used to prevent osmoadaptation. Bar graphs show mean, ± SD, n > 30. Statistical comparisons are by Student's t-test: *$p<0.05$, ***$p<0.001$, N.S., not significant.

#### Figure 4; Figure Supplement 2

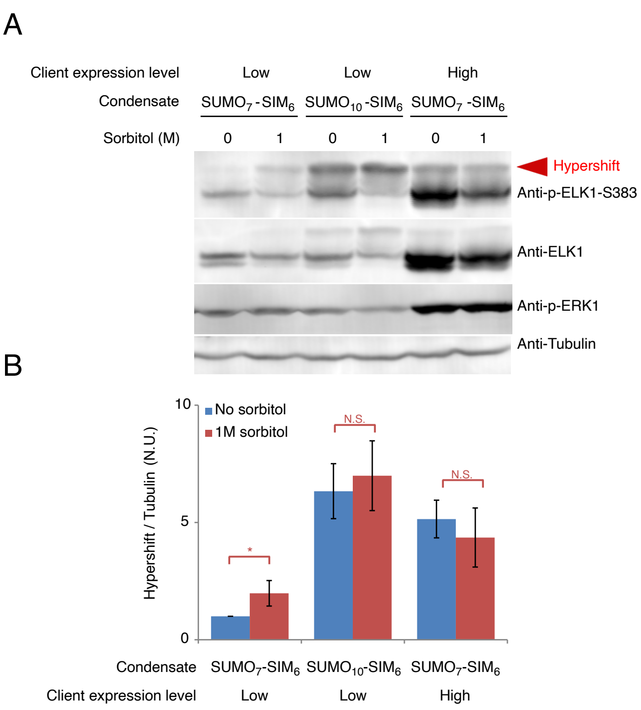


**Figure 4; Supplement 2: Phosphorylation within $SUMO_7$-$SIM_6$ condensates only responds to osmotic compression when client concentration is relatively low.** We compared ELK1 hyperphosphorylation before and after osmotic compression with 1M sorbitol for 1 hour in $SUMO_7$-$SIM_6$ condensates with low client concentration (left), $SUMO_{10}$-$SIM_6$ condensates (middle) and $SUMO_7$-$SIM_6$ condensates with high client concentration (right). Representative western blots are shown for (from top to bottom) the ELK1-S383 phosphoepitope, total ELK1, the activation-loop phosphates on the ERK1 kinase (pT202/pY204), and tubulin loading control. Quantification of the fraction of slow-migrating, hyperphosphorylated bands is quantified in bottom graphs. Blue bars are before, and red bars after, osmotic compression. Intensities of total hyperphosphorylated ELK1 (red arrow) were normalized to tubulin levels, and $SUMO_7$-$SIM_6$ condensates with low client concentration were used as a reference. *hog1Δ* strains were used to prevent osmoadaptation. Error bars indicate ± SD, n = 3, Statistical comparisons are by Student's t-test: * $p<0.05$, N.S., not significant.

#### Figure 4; Figure Supplement 3

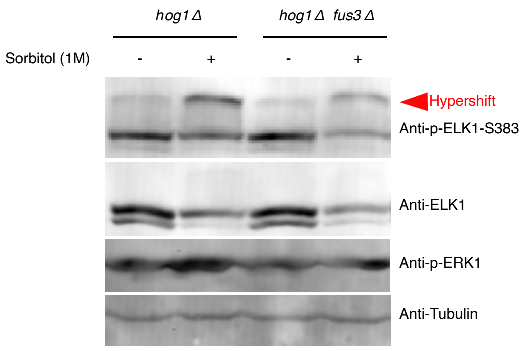


**Figure 4; Supplement 3: The Fus3 kinase is not responsible for increased hyperphosphorylation in SUMO$_7$-SIM$_6$ condensates after osmotic compression.** We compared ELK1 hyperphosphorylation before and after osmotic compression with 1M sorbitol for 1 h in SUMO$_7$-SIM$_6$ condensates in the presence (left, *hog1Δ*), or absence (right, *hog1Δ; fus3Δ*) of the Fus3 kinase. Representative western blots are shown for (from top to bottom) the ELK1-S383 phosphoepitope, total ELK1, the activation-loop phosphates on the ERK1 kinase (pT202/pY204), and tubulin loading control. Red arrowheads indicate hypershifted bands.

#### Figure 4; Figure Supplement 4

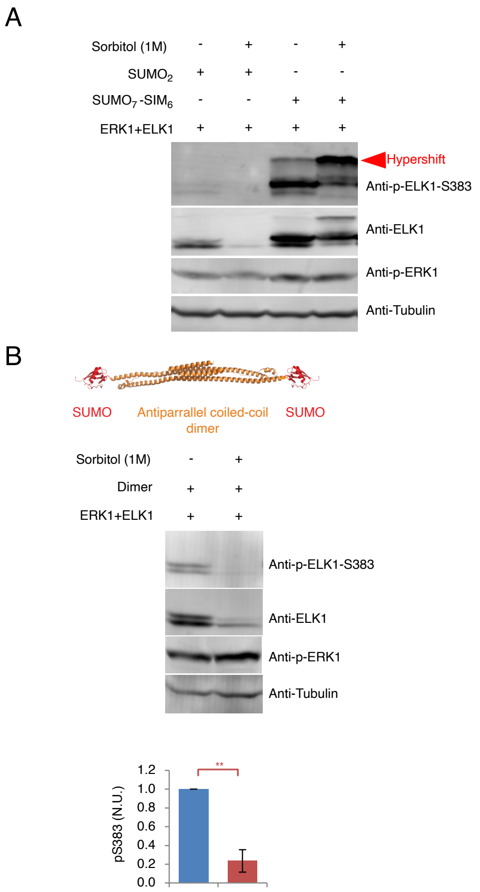


**Figure 4; Supplement 4: Osmotic compression decreases phosphorylation rates and leads to ELK1 degradation in the solute phase.** (A) Western blots comparing the effects of osmotic compression on ELK1 phosphorylation scaffolded by a simple $SUMO_2$ scaffold in the solute phase (left) versus phosphorylation in $SUMO_7$-$SIM_6$ condensates (right). Representative western blots are shown for (from top to bottom) the ELK1-S383 phosphoepitope, total ELK1, the activation-loop phosphates on the ERK1 kinase (pT202/pY204), and tubulin loading control. Red arrowheads indicate hypershifted bands. (B) Osmotic compression decreases phosphorylation rates and leads to ELK1 degradation in the solute phase in strains expressing two SUMO domains connected by a rigid coiled-coil dimer (the dimer component of the two-component system alone). Representative western blots are shown for (from top to bottom) the ELK1-S383 phosphoepitope, total ELK1, the activation-loop phosphates on the ERK1 kinase (pT202/pY204), and tubulin loading control. Hypershifted bands could not be detected in this strain, therefore total S383 phosphorylation was quantified and shown below. Band intensities of total phosphorylated ELK1 were normalized to total ELK1 levels, and this value was further normalized to the phosphorylation level of ELK1 in the control strain (left) without sorbitol treatment. *hog1Δ* strains were used to prevent osmoadaptation. Error bars indicate ± SD, n = 3, Statistical comparisons are by Student's t-test: ** $p < 0.01$, N.S., not significant.

#### Figure 4; Figure Supplement 5

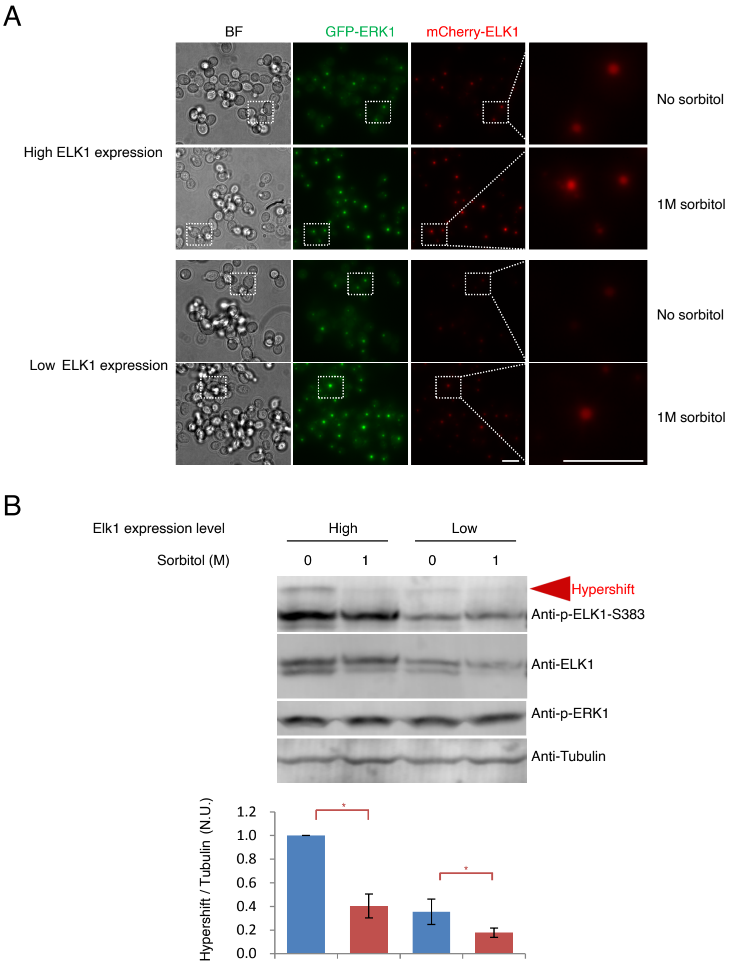


**Figure 4; Supplement 5: Osmotic compression reduces ELK1 hyperphosphorylation in synthetic two components condensates, even with lower ELK1 expression levels.**

(A) Similar increases in client concentration in two-component condensates were observed after 1 hour of osmotic compression with 1M sorbitol treatment in strains expressing low (top) and high (bottom) levels of ELK1. GFP and mCherry images represent average-intensity z-projections of 4.8 µm z-stacks with 0.4 µm step-size. Scale bar = 5 µm. (B) We compared the effect of 1 hour of osmotic compression with 1M sorbitol on ELK1 hyperphosphorylation in two-component condensates in strains with high and low ELK1 expression levels. Representative western blots are shown for (from top to bottom) the ELK1-S383 phosphoepitope, total ELK1, the activation-loop phosphate on the ERK1 kinase (pT202/pY204), and tubulin loading control. Red arrowheads indicate hypershifted bands that were quantified in the graph below. The intensity of hypershifted ELK1 bands were normalized to tubulin expression. The hyperphosphorylation level of ELK1 in the leftmost strain was set as 1. *hog1Δ* strains were used to prevent osmoadaptation. Error bars indicate ± SD, n = 3. Statistical comparisons are by Student's t-test: * $p<0.05$.

#### Figure 4; Figure Supplement 6

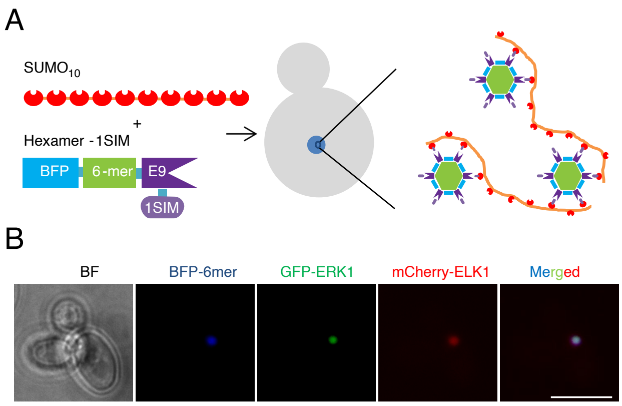


**Figure 4; Supplement 6: Co-expression $SUMO_{10}$ and a hexamer-SIM component creates a mixed condensate that can recruit SIM-tagged clients.**

(A) Schematic showing the design of a mixed system with moderate flexibility. One component consists of a homohexamerizing scaffold fused to the blue fluorescent protein (BFP), E9 domain and SIM tag. The other component consists of $SUMO_{10}$, in which each SUMO was connected by flexible $(GGS)_4$ linker. The multivalent interaction between $SUMO_{10}$ and SIM-tagged hexamer would assemble into a relatively flexible condensate. (B) The moderately flexible condensate assembled in cells coexpressing $SUMO_{10}$ and the SIM-tagged hexamer, and SIM-tagged clients (GFP-ERK1 and mCherry-ELK1) were recruited into the condensate. Images represent average-intensity z-projections of 4.8 µm z-stacks with 0.4 µm step-size. Scale bar = 5 µm.

#### Figure 4; Figure Supplement 7

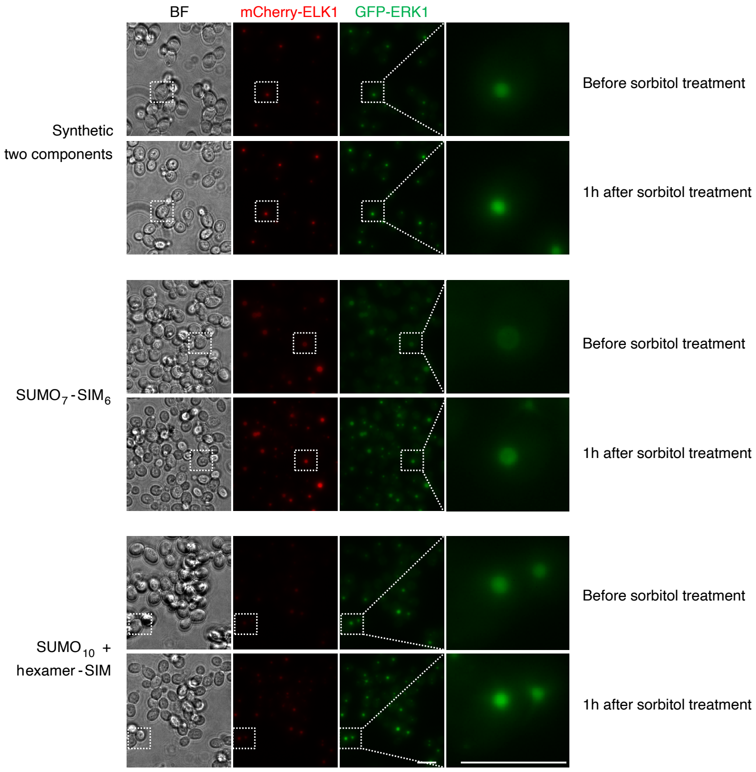


**Figure 4; Supplement 7: Osmotic compression of the moderately flexible condensate composed of the SIM-tagged hexamer and $SUMO_{10}$ had intermediate effects compared to the flexible $SUMO_7$-$SIM_6$ condensate and rigid synthetic two component system.** We compared the condensate size and client concentration in three types of condensates: the synthetic two component system (top), $SUMO_7$-$SIM_6$ (middle), and the mixed system of a SIM-tagged homohexamer (the structural scaffold of the hexamer in the two component system) with a $SUMO_{10}$ polypeptide (bottom). Cells were immobilized and imaged before and after 1 hour of osmotic compression with 1M sorbitol. Representative micrographs are shown. GFP and mCherry images represent average-intensity z-projections of 4.8 µm z-stacks with 0.4 µm step-size. Scale bars = 5 µm. Quantification of changes of client concentration and condensate area under osmotic compression is shown in Figure 4B and 4C, respectively. *hog1Δ* strains were used to prevent osmoadaptation.

#### Figure 5; Figure Supplement 1

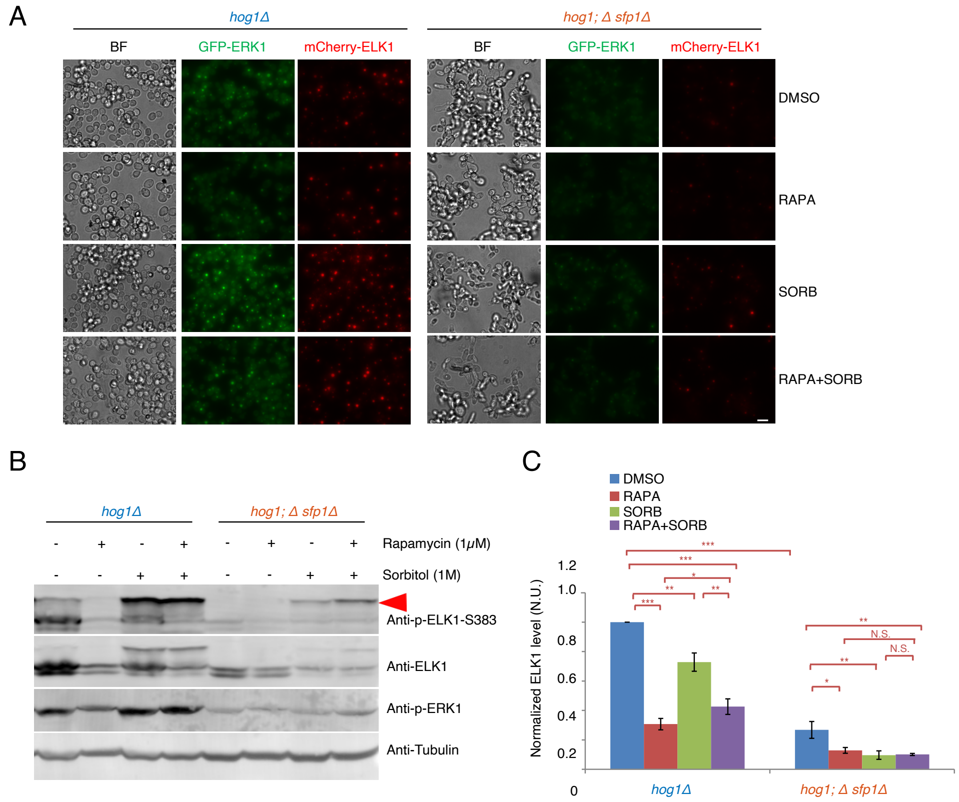


**Figure 5; Supplement 1: Reduction of client recruitment and ELK1 phosphorylation by rapamycin treatment could be reversed by sorbitol treatment**

(A - B) We compared the recruitment of GFP-ERK1-SIM in $SUMO_7$-$SIM_6$ condensates in *hog1Δ* and *hog1Δ; sfp1Δ* cells after different macromolecular crowding perturbations. Cells were treated with 1 μM rapamycin/DMSO for 2 hours, 1M sorbitol for 1 hour, or combination of both. (A) GFP and mCherry channel images represent average-intensity z-projections of 4.8 μm z-stacks with 0.4 μm step-size, scale bar = 5 μm. (B) ELK1 phosphorylation changes were evaluated after macromolecular crowding perturbations. Cells were treated with 1 μM rapamycin/DMSO for 2 hours, 1M sorbitol for 1 hour, or 1 μM rapamycin/DMSO for 1 hour followed by 1 hour treatment with both 1M sorbitol and 1 μM rapamyclin/DMSO. Representative western blots are shown for (from top to bottom) the ELK1-S383 phosphoepitope, total ELK1, the activation-loop phosphate on the ERK1 kinase (pT202/pY204), and tubulin loading control. The ELK1 phosphorylation quantification is shown in Figure 5E. The red arrowhead indicates hypershifted bands. (C) Quantification of ELK1 protein level after macromolecular crowding perturbations. Cells were treated in the same way with (B). The band intensities of ELK1 were normalized to tubulin levels, and this value was further normalized to ELK1 level in the leftmost strain (*hog1Δ* with DMSO). All bar graphs show mean ± SD, n = 3; statistical comparisons are by Student's t-test: *p < 0.05, ** p < 0.01, *** p < 0.001, N.S., not significant.

#### Figure 5; Figure Supplement 2

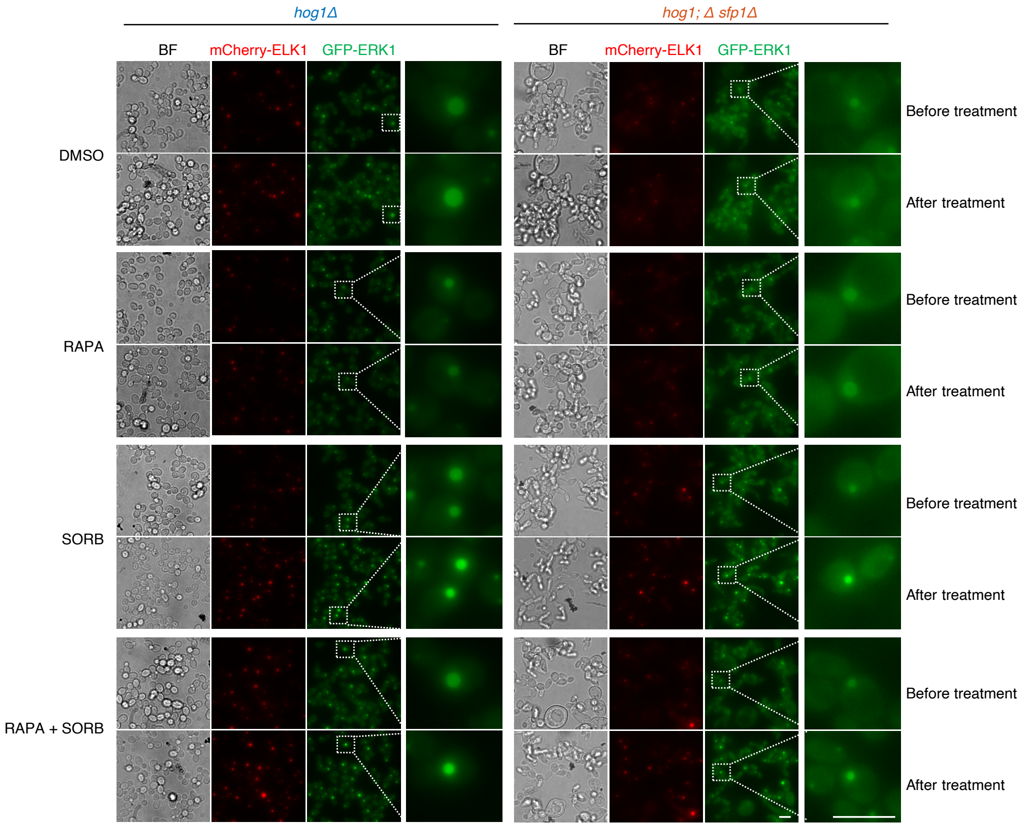


**Figure 5; Supplement 2: Macromolecular crowding perturbations alter client concentration in SUMO$_7$–SIM$_6$ condensates and SUMO$_7$–SIM$_6$ condensate area.** Cells were treated with 1 µM rapamycin/DMSO for 2 hours, 1M sorbitol for 1 hour, or 1 µM rapamycin/DMSO for 1 hour followed by 1 hour treatment with both 1M sorbitol and 1 µM rapamyclin/DMSO. Cells were immobilized and imaged before and after treatment. Representative micrographs are shown. GFP and mCherry channel images represent average-intensity z-projections of 4.8 µm z-stacks with 0.4 µm step-size. Quantification of the changes of SUMO$_7$–SIM$_6$ condensates area and client concentration in SUMO$_7$–SIM$_6$ condensates are shown in Figure 5C and 5D, respectively. Scale bar = 5 µm.

#### Figure 5; Figure Supplement 3

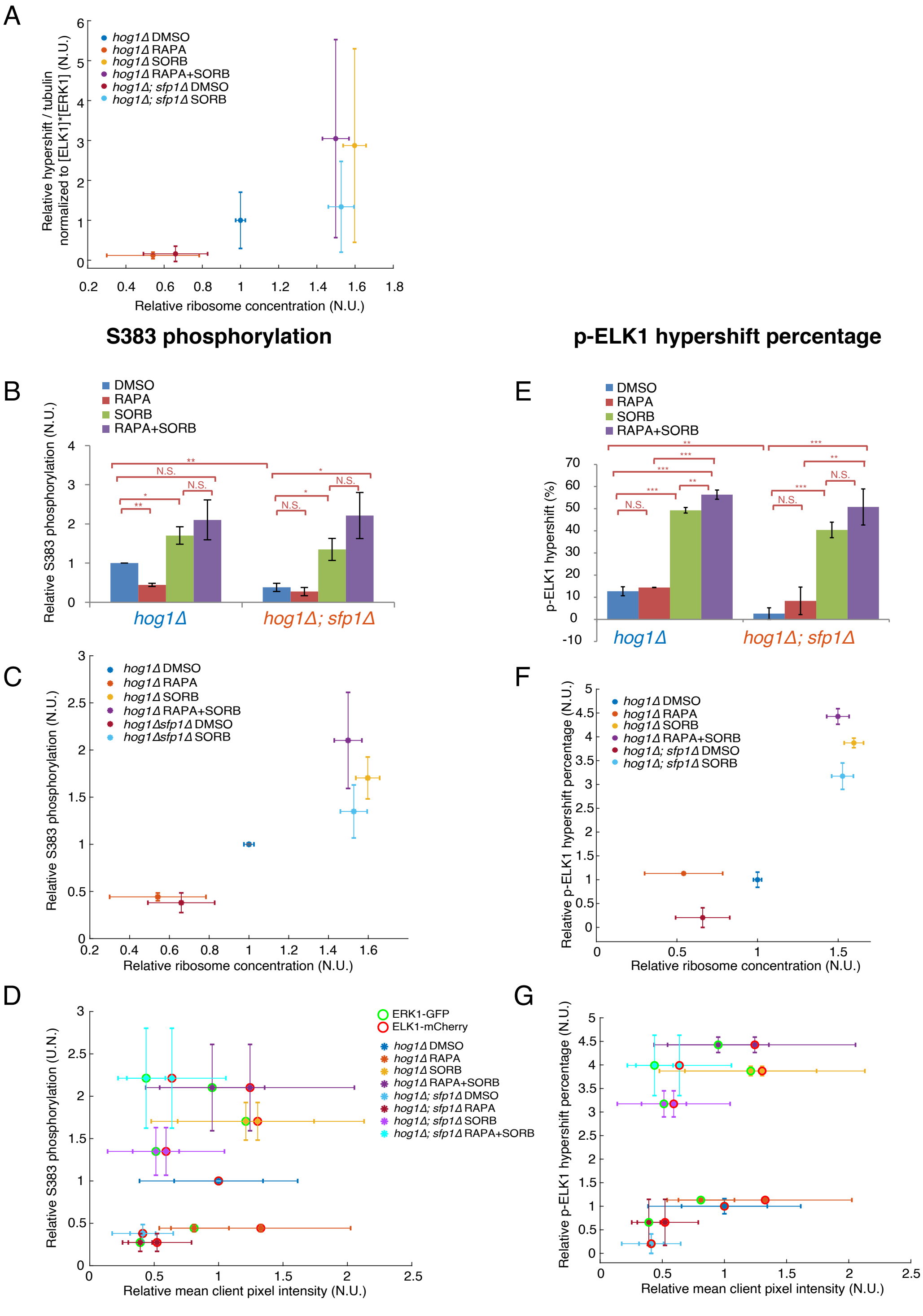


**Figure 5 - Supplement 3: Additional methods of quantifying droplet phosphorylation properties under low client concentrations.**

(A) Scatter plot of relative hypershift / tubulin (data from Fig. 5F) normalized to the product of mean GFP-Erk1-SIM and mCherry-Elk1-SIM client pixel intensity (data from Fig. 5E) under each condition were plotted against relative ribosome concentration (data from Fig. 5C). Vertical error bars are standard deviations while horizontal error bars are standard error of means. (B-

D) Ser383 phosphorylation quantification from western blot: intensities of total phosphorylated Elk1 bands (including hypershifted and non-hypershifted p-Elk bands) divided by total Elk1 band intensities (including hypershifted and non-hypershifted Elk bands). (B) Relative Ser383 phosphorylation were calculated by normalizing Ser383 phosphorylation under different conditions to Ser383 phosphorylation level in the *hog1Δ* DMSO condition. Bar graphs show mean, ± standard deviation. (C) Scatter plot of relative total Ser383 phosphorylation from B, plotted against relative ribosome concentration from Fig. 5C. Vertical error bars are standard deviations while horizontal error bars are standard error of means. (D) Scatter plot of relative Ser383 phosphorylation data from B were plotted against relative mean GFP-Erk1-SIM (green circle) and mCherry-Elk1-SIM (red circle) client pixel intensity from Fig. 5E. Error bars are standard deviations. (E-F) p-Elk hypershift percentage quantification from western blot: band intensities of hypershifted phosphorylated Elk1 were divided by total phosphorylated Elk1 bands intensities (including hypershifted and non-hypershifted p-Elk bands). (E) p-Elk hypershift percentage under different conditions were plotted using bar graphs, showing mean, ± standard deviation. (F) Relative p-Elk hypershift percentage were calculated from p-Elk hypershift percentage normalizing to its value in *hog1Δ* DMSO condition. Scatter plot of relative p-Elk hypershift percentage data were plotted against relative ribosome concentration from Fig. 5C. Vertical error bars are standard deviations while horizontal error bars are standard error of means. (G) Scatter plot of relative p-Elk hypershift percentage data were plotted against relative mean GFP-Erk1-SIM (green circle) and mCherry-Elk1-SIM (red circle) client pixel intensity from Fig. 5E. Error bars are standard deviations. Different conditions were represented through different colors of lines or dots or circle inside. All statistical comparisons are by Student's t-test: *$p<0.05$, ***$p<0.001$, N.S., not significant.

**Supplemental Movie 1 The NLS-GFP-2xWW dynamically changes localization during mitosis.** Four fields of view containing cells exiting mitosis. Log-phase cells were immobilized on a 384-well imaging plate, and z-stacks (4.8 µm stacks, 0.4 µm between focal planes) were acquired every 10 minutes. Cells were imaged using Highly inclined thin illumination (HILO) with a 488nm excitation laser and a 100x 1.4 NA oil objective. Average projections of a merge of the GFP channel (GFP-2xWW reporter, top) and the mCherry channels (mCherry-ELK1, red) are shown.

**Supplemental Movie 2 Microrheology using 40nm-GEM nanoparticles.**
Representative movies of GEM particles in *S. cerevisiae* in various conditions. Stills and time projections of tracking data from these movies are presented in Figure 5B. Cells were imaged using Highly inclined thin illumination (HILO) with a 488nm excitation laser and a 100x 1.4 NA oil objective. Each GEM movie was recorded at a single focal plane at 100 Hz (10 ms frame rate) for 4 seconds. Top, *hog1Δ* cells, bottom *hog1Δ; sfp1Δ* cells. Cells were treated with DMSO (solvent control), 1 µM rapamycin (RAPA) for 2 h to inhibit mTORC1, 1M sorbitol (SORB) or 1 µM rapamycin followed by 1M sorbitol (RAPA SORB).
